## Supporting Information for "Simulation-Driven Design of Stabilized SARS-CoV-2 Spike S2 Immunogens"

**Table of Contents**

1. **Extended Computational Methods**
    1.1. Model preparation of the HexaPro-SS-Δstalk
    1.2. Alternative simulation methods
        1.2.1. All-atom molecular dynamics (MD)
        1.2.2. Gaussian accelerated molecular dynamics (GaMD)
    1.3. Weighted ensemble (WE) simulations
    1.4. Analysis of WE simulations
    1.5. Non-equilibrium alchemical free energy calculations
2. **Supplementary Figures 1 – 25**
3. **Supplementary Tables 1 – 4**
4. **Captions for Supplementary Movies 1,2**
5. **Supplementary References**

### 1. Extended Computational Methods

#### 1.1. Model preparation of the HexaPro-SS-Δstalk

The all-atom model of the HexaPro-SS-Δstalk spike's S2 trimer, was constructed from a cryo-EM structure of the SARS-CoV-2 HexaPro spike (PDB ID: 6XKL).[1] A segment of the S2 N-terminus and the stalk residues were excluded, and only residues 696 to 1141 were included. The missing fusion peptide region (residues 823 to 857) was grafted from a cryo-EM structure of the SARS-CoV-2 3Q-2P spike (PDB ID: 7JJI).[2] As the structure of the closed prefusion HexaPro-SS-Δstalk was unavailable at the time of model preparation, the stabilizing interprotomer mutations S704C, K790C, and Q957E were incorporated with VMD's psfgen 2.0 structure-building tool.[3,4] The model was glycosylated using the same glycoprofile from Casalino et. al.[5], which represents an asymmetric, heterogenous glycan composition from glycoanalytic data.[6,7] Protonation states of the glycosylated HexaPro-SS-Δstalk model were assigned using PROPKA3[8] at pH 7.4. The model was parameterized using psfgen and the all-atom additive CHARMM36m force field for protein and glycans.[9–11] The system was solvated with TIP3P[12] explicit water molecules in a box with at least 15 Å between the protein and box edges (box size: 134 Å x 140 Å x 153 Å), and the charge was neutralized with $Na^+$ and $Cl^-$ ions up to a concentration of 150 mM. The *solvate* and *autoionize* plugins included in VMD[3] were used for solvation and charge neutralization, respectively. The final system accounted for 274,082 atoms.

#### 1.2. Alternative simulation methods.

##### 1.2.1. Molecular dynamics (MD) simulations.

All-atom conventional MD simulations were performed on the Triton Shared Computing Cluster (TSCC) at the San Diego Supercomputer Center (SDCS) using NAMD3.[13] To reduce unfavorable interactions, the HexaPro-SS-Δstalk system was energy minimized and equilibrated. An energy minimization was initially performed for 10,800 steps utilizing the conjugate gradient method. During this phase, protein and glycan atoms were subjected to positional harmonic restraints, applying a force constant of 1.0 (kcal/mol)/$Å^2$. Following this, a 0.5 ns NPT (isothermal-isobaric) equilibration was conducted, using an anisotropic pressure coupling scheme (*useFlexibleCell yes*) The harmonic restraints applied previously were retained throughout this stage. Control of the temperature and pressure was achieved with a Langevin thermostat[14] (310 K) and a Nosé-Hoover Langevin barostat[15] (1.01325 bar) as implemented in NAMD. Next, restraints were released, and 5 ns of NPT equilibration was performed upon switching to isotropic pressure coupling (*useFlexibleCell no*). Finally, 4 replicates of NPT production MD simulations were run (Rep 1 = 601.5 ns, Rep 2 = 655.4 ns, Rep 3 = 744.6 ns, Rep 4 = 627.6 ns) after re-initializing velocities. All simulations (including equilibration) were performed using an integration time step of 2 fs and the SHAKE[16] algorithm to keep covalent bonds involving all hydrogen atoms fixed. Periodic boundary conditions were active, with the particle-mesh Ewald[17] method used for long-range electrostatics calculation (every 3 timesteps). A maximum grid spacing of 2 Å was set. Non-bonded interactions, such as van der Waals interactions and short-range electrostatics, were accounted for using a cutoff distance of 12 Å.

**1.2.2. Gaussian accelerated molecular dynamics (GaMD) simulations.**

GaMD simulations were performed on the Triton Shared Computing Cluster (TSCC) at the San Diego Supercomputer Center (SDCS) using GaMD[18,19] with NAMD2.14[13]. The initial state for GaMD simulations was selected from the last frame of the first replica of all-atom conventional MD simulations, which represents a tightly closed conformation that is more compact relative to the initial state used in all-atom MD simulations. To determine the magnitude of the dual boost potential $\Delta V$ (Total potential boost + Dihedral potential boost) required to enhance conformational sampling, the reference E threshold for application of the dual harmonic boost potential was assessed at both low acceleration (E=1) and high acceleration (E=2). To obtain the optimal boost potential parameters for GaMD production simulations, an NPT equilibration was performed at 310 K and 1.01325 bar with the following protocol. The system was simulated without a boost applied for 0.4 ns without collecting system potential statistics ($V_{min}$, $V_{max}$, $V_{avg}$, $\sigma_V$), then the system was simulated for 1.6 ns where system potential statistics were collected. A boost potential is then applied for the remainder of GaMD equilibration and production simulations. Using the previously calculated system potential statistics, the system was simulated with boost potential parameters fixed for 0.4 ns, then the boost parameters were updated as the system was further equilibrated for 24.6 ns. At the end of system equilibration, the final calculated boost parameters were fixed and maintained for GaMD production simulations. Velocities were randomly initialized, and 3 replicates of NPT GaMD production simulations were run for each acceleration level at 310 K and 1.01325 bar.

### 1.3. Weighted ensemble (WE) simulations.

The weighted ensemble (WE) methodology is an enhanced sampling strategy for studying rare events in complex biological systems.[20] WE strategy involves running numerous MD simulations in parallel generating as many trajectories, also called 'walkers," of length $\tau$, each one carrying a statistically determined weight. At regular time intervals, $\tau$, a resampling process is implemented to enhance the abundance of 'promising' trajectories that have made progress toward the target state. Promising trajectories are determined based on one (or more) progress coordinates chosen between the starting and desired states (such as closed and open), which are divided into bins. A cycle of MD simulation and resampling defines a WE 'iteration.' It is important to note that the WE strategy relies on stochastic dynamics.[20] Therefore, the use of a stochastic thermostat to couple MD simulations with, such as the Langevin thermostat, where pseudo-random values are generated using a random seed based on the current date and time to re-initialize velocities before each simulation (i.e., walker), is suitable.[20] Moreover, trajectory weights are adjusted using rigorous statistical principles to ensure that no bias is introduced into the system dynamics. This implied that all the initial trajectories carry equal statistical weights, whose sum is equal to one. Then, all the trajectories are closely monitored to ensure that the sum of weights remains one throughout the simulation, thereby maintaining unbiased dynamics.[21] At each resampling step, trajectories that move to empty bins are replicated, and their weights are evenly distributed among the resulting offspring trajectories. Trajectories that fail to make progress are occasionally terminated, and their weights are combined with other trajectories that will be continued. This strategy allows for the computation of rate constants for reaching any desired state, provided that enough sampling has been carried out. WE simulations can be performed under non-equilibrium steady-state or equilibrium conditions.[22,23] At odds with non-equilibrium steady-state WE simulations, equilibrium WE simulations do not require the definition of a target state in advance. Moreover, in non-equilibrium steady-state WE simulations trajectories reaching the target state are 'recycled' back to the initial state with the same weights to ensure that non-equilibrium steady-state conditions are maintained.[22,23] In this work, we performed equilibrium WE simulations to investigate the opening mechanism of the HexaPro-SS-Δstalk construct.

*Preparation of the initial HexaPro-SS-Δstalk construct for WE simulations.*

The initial HexaPro-SS-Δstalk construct for WE simulations was generated as follows. Previous MD simulations of the HexaPro-SS-Δstalk performed with NAMD3 using CHARMM36m all-atom additive force fields[9–11] (see above for details) were parsed to extract a frame in which the three P987 residues atop the central helices (CH) formed an imaginary equilateral triangle where the distances between respective Cα atoms were as close as possible to a value of ~20 Å, as in the reference, "closed" structure with PDB ID: 6XKL.[1] A frame after 72.3 ns of the first MD replica previously performed with NAMD3 was identified as the best match, with the three $P987_{C\alpha}$–$P987_{C\alpha}$ being 18.7 Å, 18.2 Å, and 18.8 Å, respectively. The CHARMM36m topological information and the coordinates of the solute (protein and glycan atoms) at the selected frame were extracted using VMD.[3] The dry construct was re-solvated with TIP3P[24] water molecules in an orthorhombic box of 138 Å x 150 Å x 168 Å per side, allowing at least 24 Å between the solute and the box edges in all directions, and neutralized with a 150 mM concentration of NaCl using the *solvate* and *autoionize* plugins included in VMD, respectively.[3] As a result, we obtained a system tallying 328,200 atoms. In order to use the Amber software[25] as the engine for the subsequent MD simulations while maintaining the CHARMM36m force fields, we used the CHAMBER[26] program to convert the generated coordinate (*.pdb*) and topology (*.psf*) files, and the necessary CHARMM36m parameters files into an Amber readable format (i.e., *.prmtop* file for topology and parameters and *.rst7* file for coordinates).

*Generation of initial basis states for WE simulations.*

In order to ensure a more extensive sampling of the initial state in a WE simulation, it is recommended to provide a set of initial, equally weighted conformations (called "basis states") rather than a single conformation. In this work, we generated 50 initial basis states, all in the closed conformation, to which we assigned an initial weight of 0.02 each. To accomplish this task, we performed 10 preparatory MD simulations of 50 ns each of the Amber-suitable HexaPro-SS-Δstalk solvated system generated as described in the previous section, yielding a total aggregate sampling of 500 ns. This large conformational ensemble was used to extract the 50 basis states. MD simulations were performed on the UC San Diego's TSCC using Amber20 software[25,27] and CHARMM36m force fields,[9–11] adopting the following protocol for each of the 10 replicas. An initial minimization of the construct was carried out in two subsequent stages. In the first stage, only the water molecules and ions were allowed to move, while imposing a harmonic restraint of 100 (kcal/mol)/Å$^2$ to protein and glycan atoms. The restraints were lifted in the second stage, and all the atoms were allowed to move. A maximum number of 100,000 minimization steps was set in both stages, where the initial 10 steps were performed using the steepest descent method, whereas the necessary remaining steps (until convergence) with the conjugate gradient method. Next, we conducted 300 ps of heating in the NVT ensemble, where the temperature of the system was gradually brought from 0 K to 300 K in six consecutive steps of 50 ps each.[28] Temperature was controlled using Amber's implementation of Langevin thermostat with a collision frequency of 1 ps$^{-1}$.[29] Pseudo-random values for the Langevin thermostat were generated using a random seed based on the current date and time (*ig=-1*). After heating, we conducted 1 ns of equilibration in NPT conditions where pressure was set to 1 bar and controlled with isotropic scaling using Amber's implementation of Monte Carlo barostat.[30] Volume change attempt was performed every 100 steps, whereas the pressure relaxation time was set to 1 ps. For each replica, production simulation was performed for 50 ns in the NPT ensemble using a time-step of 2 fs. The temperature was maintained at 300 K (Langevin thermostat[29]), and pressure was kept at 1 bar (Monte Carlo barostat[30]). Hydrogen-containing bonds were constrained using the SHAKE algorithm.[16] Periodic boundary conditions were implemented in all simulations, and long-range electrostatic interactions were handled using the Particle Mesh Ewald method.[17] A cutoff of 10 Å was set for the calculation of non-bonded van der Waals interactions and short-range electrostatic interactions. Frames were saved every 20 ps. An aggregate of 25,000 frames was collected across the 10 replicas, corresponding to 500 ns of sampling.

For the selection of the basis states for WE simulations, we discarded the first 5 ns of production simulation for each replica. Next, we considered only the frames where the imaginary apical triangle formed by joining the three $P987_{C\alpha}$ atoms (one for each protomer) located atop the S2's central helices was equilateral, i.e., the integer part of the three $P987_{C\alpha}$–$P987_{C\alpha}$ distance values was identical. Amongst this subset of frames, we selected as initial basis states for WE simulations the 50 conformations with the largest values of $P987_{C\alpha}$–$P987_{C\alpha}$ distance, as long as the distance was larger than 15 Å and the frames separated by at least 200 ps from each other if coming from the same replica. We note that in all the 50 selected conformations, the $P987_{C\alpha}$–$P987_{C\alpha}$ distance never exceeded 20 Å.

*WE simulations of HexaPro-SS-Δstalk construct.*

WE MD simulations were performed on TSCC facility's RTX 3090 GPU nodes using the open-source, highly scalable WESTPA 2.0 software,[31] exploiting Amber20 PMEMD.cuda[25,27] as MD engine to propagate dynamics. CHARMM36m force fields[9–11] were adopted for protein and glycan, and TIP3P[24] for water molecules. WE simulations were initiated from 50 basis states in the closed conformation, each carrying an initial statistical weight of 0.02. To enhance sampling of closed-to-open transitions of HexaPro-SS-Δstalk trimer we defined two progress coordinates. The first progress coordinate, hereafter also referred to as "$P987_{TRIANGLE-AREA}$," was set as the area of the triangle having the Cα atom of residues $P987_A$, $P987_B$ and $P987_C$ as vertices. The second progress coordinate, hereafter also referred to as "$RMSD_{CH}$" was set as the Root Mean Squared Deviation (RMSD) of the Cα atoms of the CH (residues 987-1033) in the simulation construct to the corresponding Cα atoms of the CH (residues 987-1033) in the HexaPro-SS-Δstalk open crystal structure (PDB ID: 8U1G[32]). The minimal adaptive binning (MAB) scheme was applied to bin the conformational space defined by the two progress coordinates, with 15 MAB linear bins placed per progress coordinate plus two bins for the extrema and bottleneck trajectories, respectively.[33] The resampling time interval $\tau$ was set to 100 ps, whereas the target number of independent trajectories run in each bin, $M$, was set to 8. The value of the two reaction coordinates at each resampling time interval was calculated with CPPTRAJ.[34] MD simulations were performed in NPT conditions using a Langevin thermostat[29] to maintain the temperature at 300 K and the Monte Carlo barostat[30] to keep the pressure at 1 bar. SHAKE algorithm was used to constrain bonds involving hydrogen atoms.[16] Particle Mesh Ewald method was used to handle long-range electrostatic interactions,[17] whereas non-bonded van der Waals interactions and short-range electrostatic interactions were calculated with a cutoff of 10 Å.

WE simulations of HexaPro-SS-Δstalk were initially carried out for 200 iterations, resulting in ~26.9 μs aggregate sampling and 120 successful opening pathways (see Supplementary Table 3). In order to be able to provide an estimate of kinetic rates for the opening transition, we further extended WE simulations by 114 iterations, achieving a total of 315 iterations, ~46 μs aggregate sampling, and 180 successful opening pathways (Supplementary Fig. 12 and Supplementary Table 3). To facilitate the convergence of WE simulations toward equilibrium, we applied the WE Equilibrium Dynamics (WEED) protocol twice to reweight the trajectories,[35,36] specifically at the end of iterations 201 and 261. For this purpose, we used the

WEED plugin of WESTPA 2.0, setting in both instances a window size of 0.75 and dividing the conformational space defined by the two progress coordinates in 120 fixed bins (15 along the first progress coordinate, i.e., the $P987_{C\alpha}$ triangle area, and 8 along the second progress coordinate, i.e., the $RMSD_{CH}$ to the $CH_{Open\text{-}crystal}$). We note that this procedure utilizes the trajectories that are generated during the ordinary WE simulations to determine the conditional probabilities ($k_{ij}$) to hop among bins, i.e., the inter-bin rates $k_{ij}$. Then, these bin-to-bin transition probabilities are used to estimate the new bins' weights to infer steady-state behavior.[35,36]

*Modeling and simulation setup of HexaPro-SS-V991W, HexaPro-SS-T998W, and HexaPro-SS-2W variants.*

The three HexaPro-SS-Δstalk tryptophan-stabilized systems simulated in this work (HexaPro-SS-V991W, HexaPro-SS-T998W, and HexaPro-SS-2W) were built from the same base construct used for the preparation of the HexaPro-SS-Δstalk WE simulations, i.e., a frame generated after 72.3 ns of the initial MD simulations of HexaPro-SS-Δstalk performed with NAMD3. See the *Preparation of the initial HexaPro-SS-Δstalk construct for WE simulations* section above for additional information on the base construct. To generate the HexaPro-SS-V991W mutant system, we incorporated the V991W mutation on each protomer using the psfgen 2.0 structure-building tool within VMD using the *mutate* command.[3,4] In a similar fashion, we built the HexaPro-SS-T998W mutant system by incorporating the T998W mutation on each protomer, whereas for the HexaPro-SS-2W mutant system, we introduced both V991W and T998W mutations on each protomer. The mutated residues and the residues located within 5 Å of them were locally minimized for 1000 steps using the AutoIMD tool[37] within VMD.[3] Subsequently, the three dry mutated constructs were solvated using the VMD's *solvate* plugin[3] by adding TIP3P[24] water molecules to create an orthorhombic box measuring 138 Å x 150 Å x 168 Å on each side. The solute was positioned at least 24 Å away from the box edges in all directions. The system was neutralized with a 150 mM concentration of NaCl using the *autoionize* plugin within VMD.[3] The three final constructs tallied 328,328 (HexaPro-SS-V991W), 328,271 (HexaPro-SS-T998W), and 328,370 (HexaPro-SS-2W) atoms. Similarly to what was done for the HexaPro-SS-Δstalk base construct, we used the CHAMBER[26] program to convert the generated coordinate (.pdb) and topology (.psf) files into a format compatible with Amber software[25] (specifically, a .prmtop file for topology and parameters, and an .rst7 file for coordinates) using the necessary CHARMM36m parameter files.

In preparation for the WE simulations, we performed 10 preparatory MD simulations of 50 ns each for each solvated mutant system, yielding a total aggregate sampling of 500 ns. From this conformational ensemble, we extracted 50 closed conformations that we used as starting basis states in the respective WE simulation. For each of the 10 replicas carried out for each mutant, MD simulations were performed on the SDSC TSCC cluster using Amber20 software[25,27] and CHARMM36m force fields.[9–11] Criteria for the selection of the 50 initial basis states and simulation protocol were identical to those adopted for the HexaPro-SS-Δstalk base system. Details are described in the respective section above titled *Generation of initial basis states for WE simulations.*

*WE simulations of HexaPro-SS-V991W, HexaPro-SS-T998W, and HexaPro-SS-2W variants.*

Similar to the HexaPro-SS-Δstalk base construct, we performed WE MD simulations of HexaPro-SS-V991W, HexaPro-SS-T998W, and HexaPro-SS-2W systems on the TSCC's RTX 3090 GPU nodes using the WESTPA 2.0 software[31] in combination with Amber20 PMEMD.cuda MD engine[25,27] to propagate dynamics, CHARMM36m force fields[9–11] for protein and glycan, and TIP3P[24] for water molecules. As described above, WE simulations for the three mutants were started from 50 basis states in the closed conformation, each carrying an initial statistical weight of 0.02. WE simulations were carried out using the same progress coordinates and settings detailed above for the HexaPro-SS-Δstalk base construct in the *WE simulations of HexaPro-SS-Δstalk construct* section.

WE simulations of the HexaPro-SS-V991W mutant were initially carried out for 200 iterations resulting in ~23.5 μs aggregate sampling and 100 successful pathways. To provide a more accurate estimate of kinetic rates, we further extended WE simulations by 101 iterations, achieving a total of 301 iterations, ~35.7 μs aggregate sampling, and 149 successful opening pathways (Supplementary Fig. 12 and Supplementary Table 3). At the end of iterations 201 and 261 we reweighted the trajectories using WEED plugin of WESTPA 2.0 to improve convergence toward equilibrium.[35,36] For this purpose, the same settings detailed above for the HexaPro-SS-Δstalk base construct were used.

WE simulation of the HexaPro-SS-T998W mutant was performed for a first set of 254 iterations resulting in ~32.5 μs aggregate sampling and 155 successful opening pathways. Then, WE simulations were continued for additional 101 iterations, reaching a total of 355 iterations, ~47.7 μs aggregate sampling, and 187 successful opening pathways (Supplementary Fig. 12 and Supplementary Table 3). Trajectories were reweighted at the end of iterations 255 and 315 using WEED plugin of WESTPA 2.0.[35,36] For this purpose, the same settings detailed above for the HexaPro-SS-Δstalk base construct were used.

WE simulation of the double HexaPro-SS-2W mutant was initially executed for 269 iterations resulting in ~35.4 μs aggregate sampling and 136 successful opening pathways. We further extended the WE simulation for additional 132 iterations, reaching a total of 401 iterations, ~56.6 μs aggregate sampling, and 195 successful opening pathways (Supplementary Fig. 12 and Supplementary Table 3). Reweighted of trajectories was conducted twice, namely at the end of iterations 269 and 330 using WEED plugin of WESTPA 2.0.[35,36] For this purpose, the same settings detailed above for the HexaPro-SS-Δstalk base construct were used.

*Selection of WE progress coordinates.*

The choice of progress coordinates is critical and should directly correlate to the mechanism being investigated. To capture significant conformational changes for the HexaPro-SS-Δstalk, categorization of the states is a necessary and required step. Here, we demonstrate the conformational transition of the system from a "closed" state to an "open" state. One geometric measure of such a significant change could be the volume occupied by the system in the "closed" condition compared to the "open" state. Since we wanted the measure to be sensitive to the state change, a two-dimensional projection of the volume, i.e., the surface area, stood out as a potential progress coordinate. The area of the triangle spanned out between the α-carbon atoms of the three P987 residues of the three monomers ($P987_{TRIANGLE\text{-}AREA}$) was, therefore, an automatic choice to demonstrate the opening and closing mechanism of the system. Conformational stability and flexibility of central helices are crucial for the overall dynamics of the spike protein. RMSD is a highly relevant progress coordinate that directly measures the conformational deviations. RMSD of the three CH (residues 987-1033) with respect to CH in the open crystal structure is considered the second progress coordinate ($RMSD_{CH}$) for observing the opening transition of the S2 trimer in the WE simulations. Both the progress coordinates provide easy-to-interpret and intuitive insights into the dynamics of the S2 trimer. While $P987_{TRIANGLE-AREA}$ provides a geometric visualization of the closing and opening mechanism, $RMSD_{CH}$ highlights the magnitude of the deviation of the CH with respect to the open state, providing a comprehensive view of the protein dynamics. $RMSD_{CH}$ provided insights into the significance of these helices in the dynamics of the system, playing a pivotal role in the conformational change. RMSD profile observed in the unbiased MD simulations, in contrast to GaMD, are more dynamic, underscoring the importance of this metric in capturing the holistic and nuanced behavior of the system (Supplementary Fig. 1).

#### 1.4. Analysis of WE simulations.

*WE simulation summary.*

A summary of the weighted ensemble trajectory data for each system, including iteration number, number of walkers per iteration, minimum and maximum bin probabilities, minimum and maximum segment probabilities, cpu-time, wall-time, was generated by accessing the respective simulation data file (*west.h5*) using the *Run* interface of the *westpa.analysis* Python API.[38] Summary reports for each WE simulation are included in the Supplementary Data 1.

Convergence of WE simulations was assessed in two ways. The first one was through the 1D probability plots reported in Supplementary Figs. 2b,8b,9b,10b. For each progress coordinate, we calculated the probability distribution at three different times. Notably, a significant shift is observed between the probabilities calculated over the WE iterations preceding reweighting and those computed during the WE iterations spanning the period between the 1$^{st}$ and 2$^{nd}$ reweighting. Subsequently, after the 2$^{nd}$ reweighting, the probabilities tend to stabilize, showing only minor deviations from the previous values in the region of interest. This trend underscores reasonable convergence. Otherwise, we would have observed another large shift in the probabilities. The second way we assessed convergence is illustrated in the plots reported in Supplementary Fig. 15, where the evolution of the rolling average of the kinetic rate estimated for the opening transition is presented for each system. In a scenario where the simulations are fully converged, the curve indicating the evolution of kinetic rate rolling average values over the WE iterations should plateau around a specific value. As it can be evinced from Supplementary Fig. 15, especially from panel C, where the rolling averages were calculated over larger windows of iterations (window width corresponding to ~10% of the total number of WE iterations instead of one iteration only), for all systems the curve start plateauing, especially over the iterations following the 2$^{nd}$ reweighting.

Amongst the four systems, the convergence of WE simulations for HexaPro-SS-2W has room for improvement. However, considering the complexity of these simulations and the very good relative agreement amongst the four systems, the observed trends are very reassuring in this sense.

*Probability distribution of the progress coordinates.*

The average probability distribution as a function of the two progress coordinates was calculated for all the WE simulations using *w_pdist* and *plothist* tools included in the WESTPA 2.0 release. The *average* plotting mode was used with *plothist*, which plots the probability as a time average over a specified number of iterations. Three 2-D (considering both progress coordinates) probability distribution histograms were constructed for each system, i.e., one averaged over the WE iterations occurring before the first reweighting, the second averaged over the WE iterations occurring between the first and the second reweighting, and the third one averaged over the WE iterations performed after the second reweighting (Supplementary Figs. 2a,8a,9a,10a). The following ranges of iterations were considered to perform this analysis: *HexaPro-SS-Δstalk*: 1–201 / 202–261 / 262–315; *HexaPro-SS-V991W*: 1–201 / 202–261 / 262–301; *HexaPro-SS-T998W*: 1–255 / 256–315 / 316–355; *HexaPro-SS-2W*: 1–269 / 270–330 / 331–401. Using the same three iteration ranges and the same *average* plotting mode, three 1-D probability profiles were also produced for each progress coordinate and for each system (Supplementary Figs. 2b,8b,9b,10b). The three 1-D probability profiles obtained for each progress coordinate were incorporated into the same plot to simplify visualization. Finally, we also monitored the time evolution of the probability of each progress coordinate over the specified range of iterations using the *evolution* plotting mode of *plothist* (Supplementary Figs. 2b,8b,9b,10b, central and right panels).

*Definition of "closed" and "open" states.*

The states used for the kinetics analysis were defined in the *west.cfg* configuration file, which contains the adopted binning scheme of the two WE progress coordinates to evaluate the state membership of the conformations sampled during WE simulations. Two states were defined, namely "closed" and "open." Specifically, the "closed" state was assigned to any conformation with $P987_{TRIANGLE\text{-}AREA} \leq 200$ Å$^2$ and 9 Å $\leq RMSD_{CH} \leq$ 11 Å. The "open state" was instead assigned to any conformation with $P987_{TRIANGLE\text{-}AREA} \geq 500$ Å$^2$ and $RMSD_{CH} \leq 2$ Å. The "open" state was regarded as the target state in all the analyses performed unless otherwise specified. We defined a successful opening event as the transition between the "closed" and the "open" state (Supplementary Figs. 2b,8b,9b,10b).

*Analysis of kinetics for the opening transition from WE simulations.*

To estimate the rate constants of a transition between two separate states in an equilibrium WE simulation, the set of trajectories representing the equilibrium behavior can be partitioned into two steady states, i.e., the forward steady state and the reverse steady state.[23,35] In our work, the forward steady state is labeled as the *opening* steady state and accounts for the trajectories that ended up in the open state while having more recently visited the closed state. On the other hand, the reverse steady (not investigated here), referred to as the *closing* steady state, consists of trajectories that arrived in the closed state while having more recently been in the open state. By separately analyzing these steady states in their corresponding directions, one can estimate the rate constants from the flux probability of the trajectories arriving in the respective target state.[35] The unimolecular rate constant for the *opening* transition, $k_{opening}$, was calculated using the following equation:[23,35]

$$k_{opening} = \frac{Flux(C \rightarrow O|opening)}{p_C^{opening}} = \frac{1}{MFPT(C \rightarrow O)}$$

where $Flux(C \rightarrow O|opening)$ is the conditional flux of probability carried by trajectories in the *opening* steady state, given that the trajectories originated in the closed state ($C$ ) and arrived in the open state ($O$) at any point in the simulation; $p_C^{opening}$ is the steady state population of the closed state $C$ for the opening steady state, *i.e.* the sum of the statistical weights of trajectories more recently in the closed state $C$ than in the open state $O$.[23,35] According to the Hill relation the conditional flux of probability carried by trajectories in the *opening* steady state, corresponds to the inverse of the mean first-passage time ($MFPT(C \rightarrow O)$) for the opening transition. Normalizing $Flux(C \rightarrow O|opening)$ by the $p_C^{opening}$, i.e., the population of the closed state in the *opening* steady state ensures that the rate constant is calculated unidirectionally only for the forward *opening* steady state.[36] We note that the present work aimed to enhance the sampling of opening trajectories and pinpoint possible differences across the four mutants in the kinetics of the opening transition.

For this reason, and also because no closing successful pathways were observed, the *closing* steady state was not investigated in our analyses.

To retrieve all the successful opening events and the conditional flux probabilities of *opening* events, we used the *w_ipa* tool of WESTPA 2.0 with the *analysis-only* flag. By accessing the WE data stored in the *west.h5*, this tool allowed us to assign conformations to the "closed" and the "open" state, assess their population, and calculate the conditional probability flux of trajectories arriving in the "open" state. The analysis scheme, including state definitions (see previous section), was specified in the configuration file (*west.cfg*). The following analysis options were used. For every system, the analysis was performed considering either the entire WE simulation data (i.e., all the iterations starting from the first one) or only the WE iterations following reweighting using the default *evolution: cumulative* option to ensure that all the evolution datasets, including the conditional fluxes, were evaluated as a rolling average calculated over 1-iteration windows (*step_iter: 1*) with 95% confidence interval (*alpha: 0.05*) (Supplementary Fig. 15a,b,d). To inquire about convergence, rolling averages were also calculated over windows with a width greater than 1 iteration (Supplementary Fig. 15c), namely 10% of the total number of WE iterations (31, 40, 35, and 30 for HexaPro-SS-Δstalk, HexaPro-SS-2W, HexaPro-SS-T998W, and HexaPro-SS-V991W systems, respectively).[38] Uncertainties in the conditional probability fluxes were estimated directly with the Monte Carlo bootstrapping approach implemented within *w_ipa* tool.[21,23,39]

Results from the analysis with *w_ipa* were stored in the *assign.h5* and *direct.h5* files, the former containing state populations and the latter containing conditional probability fluxes, normalized conditional probability fluxes (i.e., kinetic rates), and event durations. The rolling average values of the conditional flux probability of the trajectories arriving in the open state, already normalized by the population of the closed state in the *opening* steady state ($p_C^{opening}$) were extracted at each iteration from the *rate_evolution/expected* dataset contained in the *direct.h5* file. These values correspond to the rate constants of the opening transition in units of inverse $\tau$. Similarly, uncertainties of rate constants (i.e., standard errors of the mean calculated with the Monte Carlo bootstrapping approach) were extracted from the *rate_evolution/sterr* dataset stored in the *direct.h5* file. Final rolling values of rate constants and relative standard errors of the mean were subsequently converted to units of inverse time ($s^{-1}$) by dividing them with the resampling time used in the WE simulations ($\tau$ = 0.1 ns = 100 × $10^{-12}$ s) (Supplementary Table 3). Considering that reweighting of WE trajectories improved convergence toward the *opening* steady state, only the estimates of the rate constants calculated over the iterations following reweighting were used. However, estimates of the rate constant employing the entire WE data, hence also including the iterations prior to reweighting, were retrieved and plotted for sake of completeness (Supplementary Table 3).

*Aggregate sampling to the first opening event and molecular time to opening.*

The total aggregate sampling until the first opening event was retrieved from the total number of completed walkers until the system reached the open state for the first time during the WE simulation (Supplementary Fig. 13a). By summing all the walkers within every iteration preceding (and including) the walker in which the opening event was observed for the first time and multiplying this number by the actual walker's length ($\tau$) used in the simulations, i.e., 0.1 ns, we obtained the total aggregate sampling value. The complete list of the number of walkers per iteration was retrieved from the *west.h5* file produced during the WE simulation.

Molecular time to opening (Supplementary Fig. 13b), i.e., the molecular time required to reach the open state, including the "waiting" time spent in the closed state, was calculated for each opening pathway by multiplying the iteration number where the system reached the open state by the resampling time ($\tau$), 0.1 ns. All the values were then represented as a distribution using a kernel density distribution plot (Supplementary Fig. 13b).

*Opening event duration.*

Durations of every opening event, i.e., the molecular time to reach the open state upon leaving the closed state, were extracted in units of $\tau$ from the *durations/duration* dataset stored in the *direct.h5* file. Durations were converted to simulation molecular times (ns) by multiplying $\tau$ with the actual value of $\tau$ used in the simulations, i.e., 0.1 ns. For each system, we calculated the distribution of opening event durations by building a histogram with bins of width 15 $\tau$ (1.5 ns) and accounting for the respective weight of each opening trajectory (Supplementary Fig. 14). The mean and the standard deviation of the mean of opening event durations were then calculated (Supplementary Table 3).

*Interprotomer distance.*

For each system, interprotomer distances were calculated for all the successful opening trajectories retrieved from the respective WE simulation. The analysis was performed with MDAnalysis.[40] At every frame of each opening trajectory, three distances were computed, namely the distance between the COM of protomer A and the COM of protomer B, the COM of protomer A and the COM of protomer C, and between the COM of protomer B and the COM of protomer C. All the Cα atoms of each protomer were considered for the estimation of the respective COM. Mean and corresponding standard deviation values of the interprotomer distances were plotted in the background of the conformational space defined by the two progress coordinates using a 2D hexagonal binning plot produced with *pyplot.hexbin* function included in the Python's Matplotlib library,[41] where the average and standard deviation values were computed using all the interprotomer distances located within each hexagon (Supplementary Fig. 12).[41] A *gridsize* value of 20 was chosen for the hexbin plot.

*Contact map analysis of the opening pathways.*

PyContact[42] was used to analyze non-covalent helix–helix contacts at each chain-chain interface of the S2 trimer. For each system, frames from the successful opening pathways were binned into closed, partially open, and open conformations based on the collective variables defined above (see *Definition of closed and open states* section); partially open conformations were assigned to frames not defined as closed or open. Helices were defined with the following residue ranges: CH (residues 978 to 1030), heptad repeat 1 helices (HR, residues 945 to 977), and upper helices (UH, residues 738 to 783). For each residue on this set of helices, we evaluated the contacts made with residues of adjacent chains (i.e., protomers) based on a distance cutoff of 5.0 Å considering side-chain atoms only. Then, by exploiting the sigmoid function implemented within the PyContact algorithm that scores contacts based on the extent of interatomic distances (i.e., higher score for tighter contacts),[43] a cumulative contact score was calculated for each residue as a sum of all the individual scores assigned to contacts made by that residue with adjacent chains' residues. As an example, the cumulative PyContact score for residue $T998_{Chain\text{-}A}$ is given by the sum of pairwise $T998_{Chain\text{-}A}$–*residue* scores for each *residue* within $CH_{Chain\text{-}B}$, $CH_{Chain\text{-}C}$, $UH_{Chain\text{-}B}$, $UH_{Chain\text{-}C}$, $HR1_{Chain\text{-}B}$, $HR1_{Chain\text{-}C}$. For additional details on how the contact score is calculated by the PyContact, refer to the original publication.[43] The cumulative contact score obtained for each residue was then normalized by the *relative contact frequency*, defined as the number of frames belonging to a given conformation with nonzero contact scores divided by all frames in successful opening pathways. We note that the contact score range for heatmap visualization was set from 0 (no contacts) to 1 (the 95$^{th}$ percentile value of the S2 base construct):

$$Contact\ score_{helix_a residue_i} = Relative\ contact\ frequency \times \sum_{a<b}^{n_{helices}} \sum_{i<j}^{n_{helix_b residues}} pycontact\ score_{helix_a residue_i - helix_b residue_j}$$

For a given state *k*, residue *i* is in helix *a,* and residue *j* is in helix *b* in a different chain.

*Contact analysis of the closed state conformations generated from WE simulations.*

To comprehensively characterize the "closed" state of each system, we systematically extracted conformations associated with this state from the entire ensemble of conformations generated during the respective WE simulation. This set encompasses even those conformations that did not lead to an actual opening event. As a criterion for selecting frames, we used the interprotomer distance between Cα atoms of P987 residues, where the maximum value for all three distances was set to 21 Å (corresponding to a $P987_{TRIANGLE\text{-}AREA}$ value of ~191 Å$^2$). A total of 50,137 frames (corresponding to 5.0137 µs) with the S2 base construct in a "closed" conformation were extracted, whereas 71,041 frames (corresponding to 7.1041 µs) for the T998W system, 44,972 frames (corresponding to 4.4972 µs) for V991W system, and 64,675 frames (corresponding to 6.4675 µs) for the V991W/T998W system.

Interprotomer contact analysis was performed with MDAnalysis.[40] The contact analysis was carried out for each system on every frame of the previously extracted dataset of closed conformations. This analysis was implemented on a pairwise residue basis using a distance cutoff of 4.5 Å and considering side-chain heavy atoms only. In this instance, a distinct tool and a stricter cutoff were used, diverging from the approach taken with PyContact for the analysis of the opening pathways (refer to the previous section). The objective here was not to score contacts based on the extent of interatomic distances but to straightforwardly gauge the frequency of contacts within the multi-µs-long conformational ensemble representative of the closed state. Residue contacts were evaluated between helices belonging to different protomers (i.e., A–B, B–C, A–C). The helices included in the analysis, for each protomer, were defined with the following residue ranges: CH (residues 978 to 1030), HR1 (residues 945 to 977), UH (residues 738 to 783). The total number of interprotomer residue contacts at each frame was retrieved by summing the number of interprotomer residue contacts between protomer pairs A–B, B–C, and A–C. The total number of interprotomer contacts involving residues at positions 991 and 998 was then filtered out. Detailed residue–residue contact information for each pair of protomers was also stored for each frame in order to calculate the relative frequency by which that specific residue–residue contact occurred throughout the analyzed dataset. We note that due to the trimeric nature of S2, for a residue–residue contact to have a relative frequency of 100%, it needs to occur 3 times at every analyzed frame, i.e., one time for each pair of protomers. Therefore, the relative frequency of a residue–residue contact was calculated as the sum of the number of times that the interaction between those two specific residues occurred across all three pairs of protomers divided by three times the total number of frames analyzed (Supplementary Figs. 19,20).

*Clustering of HexaPro-SS-2W closed state conformations.*

To extract a representative conformation of the HexaPro-SS-2W mutant in the "closed" state, we performed a K-means clustering (Supplementary Fig. 18) of the respective dataset previously generated (see previous section for details). K-Means clustering was performed using the *sklearn.cluster.KMeans* class within Python's scikit-learn package.[44] For this purpose, two features were selected, namely the $P987_{\text{TRIANGLE-AREA}}$ (see the *WE simulations of HexaPro-SS-Δstalk construct* section for details) and the number of interprotomer contacts (see previous section for details on how this was calculated). The elbow method and the analysis of the 2D kernel density of the dataset (estimated using the *kdeplot* function of Python's *seaborn*[45] visualization library) were employed to determine the appropriate number of clusters, ultimately determined as 3. Notably, while the elbow method initially indicated an optimal cluster count of 4, this number was subsequently refined to 3 upon examination of the kernel density distribution of the two features (Supplementary Fig. 18). For centroid initialization, we used the default *k-means++* method. The K-Means algorithm was allowed a maximum of 300 iterations for convergence. The resultant cluster populations were distributed as follows: 46.64% for Cluster 1, 32.64% for Cluster 2, and 20.72% for Cluster 3. Subsequently, we extracted the 50 conformations closest to each centroid from the dataset using MDAnalysis.[40] After careful visual evaluation, we chose a representative conformation from Cluster 1 for presentation in Fig. 4 of the main text.

### 1.5. Alchemical non-equilibrium free energy calculations.

*System setup.*

To build the thermodynamic cycle for free energy calculations, two independent systems representing the folded and unfolded states of the protein were generated. For the unfolded state, we used a capped tripeptide (GXG), where X is the amino acid of interest (either V, T, or W). The wild-type and the two mutated tripeptides were built with Chimera.[46] For the wild-type folded state system, we used the same equilibrated HexaPro-SS-Δstalk construct extracted from conventional MD (see *Preparation of the initial HexaPro-SS-Δstalk construct for WE simulations* section). For the constructs used in these calculations glycans were not considered; however, we note that the region of the S2 trimer surrounding the central helices does not bear glycans as the surface is buried beneath S1. Mutant systems for the folded state (HexaPro-SS-V991W, HexaPro-SS-T998W, HexaPro-SS-2W) were constructed from the wild-type system using the psfgen 2.0 structure-building tool within VMD via the *mutate* command.[3] For each variant, mutations were initially incorporated into one protomer only (protomer A), generating the respective A-mutated construct. Next, starting from this construct, a second system was built by introducing the respective mutation also in the second protomer (protomer B). Finally, a third construct incorporating the respective mutation in all three protomers (A, B, and C) was created. At each step, the mutated residues, as well as the residues located within 5 Å of them, were locally minimized for 1000 steps using the AutoIMD tool[37] within VMD.[3] The PMX[47] software was used to build hybrid structures and topologies incorporating side chains from both the wild-type and mutant proteins. Generated structures of the solute were then solvated in a cubic TIP3P[24] water box using Gromacs 2022.1 software[48,49], imposing at least 12 Å from the solute edges in each direction. $Na^+$ and $Cl^+$ ions were added at a concentration of 150 mM to neutralize the systems. Energy minimization of the generated systems was carried out using the steepest descent algorithm in Gromacs 2022.1 software.[48,49]

*Equilibrium molecular dynamics simulations.*

Generated systems were initially equilibrated in the isothermal-isobaric ensemble (NPT) for 100 ps with harmonic position restraints applied to all protein heavy atoms. Langevin dynamics[50,51] was used to couple the temperature at 298.15 K, whereas Berendsen barostat[52] was used to couple the pressure at a target pressure of 1 bar. The particle mesh Ewald algorithm[17] was employed to handle the long-range electrostatic interactions using a real space cut-off of 1 nm, a Fourier spacing of 12 Å, and a spline order of 4. The Verlet cutoff scheme was used in the neighbor list search with a cutoff of 10 Å. The P-LINCS algorithm[53] was utilized to constrain all bonds involving hydrogen atoms. All simulations were performed with CHARMM36m force fields[9,10] for protein and TIP3P[24] model for water molecules using Gromacs 2022.1 software.[48,49] Next, unrestrained equilibrium simulations were carried out for 20 ns in the NPT ensemble at a target pressure of 1 bar using the Parrinello-Rahman barostat[54] and a time step of 2 fs. The equilibration procedure was carried out independently for both wild-type and mutant systems in both the folded and unfolded states. In order to aid the convergence of the subsequent non-equilibrium MD simulations (see next section) and obtain accurate error estimates, the aforementioned procedure was repeated five times for both states (folded and unfolded) of each mutant system. Frames were saved every 100 ps, for a total of 200 frames saved per simulation.

*Alchemical non-equilibrium molecular dynamics simulations.*

200 snapshots were extracted from each preparatory unrestrained, 20 ns-long equilibrium MD simulation (one every 100 ps) for a total of 1000 snapshots for each state (folded and unfolded) considering the 5 replicas. These frames were used to spawn 1000 non-equilibrium MD simulations in the forward direction (wild-type-to-mutant) and 1000 non-equilibrium MD simulations in the reverse direction (mutant-to-wild-type) for both the folded and the unfolded state. A parameter $\lambda$ was used to control the contribution of the wild-type and mutant side chain to the Hamiltonian such that at $\lambda = 0$, the Hamiltonian corresponds to the wild-type protein, whereas at $\lambda = 1$ it corresponds to the mutant.[55] The alchemical transition between wild-type and mutant was achieved in 100,000 steps (200 ps) by switching the $\lambda$ parameter from 0 to 1 or from 1 to 0, depending on the direction of the transition, using a gradient of 0.00001/step. During these simulations, the soft-core function and the default parameters for Coulombic and van der Waals interactions were used.[56]

*Free energy calculations.*

Having multiple mutations at once imposes a strong perturbation to the system. Therefore, more work will be dissipated along the path leading to less accurate free energy estimation. To overcome this caveat, free energy calculations that include multiple mutations can be performed sequentially. Since the HexaPro-SS-Δstalk construct is a homotrimer, three subsequent free energy calculations were necessary (Supplementary Figs. 16,17). The first one was performed by introducing the mutation in chain A only. Subsequently, another independent calculation was carried out where the mutation was also introduced in chain B, with the mutation already present in chain A. Finally, an independent calculation was performed where the mutation was also induced in chain C, with the mutation already present in both chains A and B. The same workflow was implemented in the case of the double mutant (HexaPro-SS-2W), where six independent calculations were performed sequentially (Supplementary Fig. 17). For each independent calculation, PMX *analyse* script was used to extract the work values corresponding to each alchemical transition, and the free energy differences were calculated based on the Crooks fluctuation theorem[57] by utilizing the Bennett acceptance ratio (BAR) as a maximum likelihood estimator.[58] The numerical uncertainty of each calculation was estimated by employing a bootstrapping procedure (100 resampling runs for every error estimate). A thermodynamic cycle was then constructed as shown in Supplementary Fig. 16, and the difference in protein folding free energy upon mutation ($\Delta\Delta G_{folding}$) was calculated as follows:

$$\Delta\Delta G_{folding} = \Delta G_{mutated\ folded} - \Delta G_{mutated\ unfolded}$$

where the total change in free energy for the mutated trimer ($\Delta G_{mutated\ folded}$) was calculated by combining the $\Delta G$ values recovered from each independent calculation (Supplementary Table 3). *The* $\Delta G$ for the tripeptide was calculated once and multiplied by three to obtain $\Delta G_{mutated\ unfolded}$. Uncertainty propagation was applied when combining $\Delta G$ values.

### 2. Supplementary Figures 1 – 23

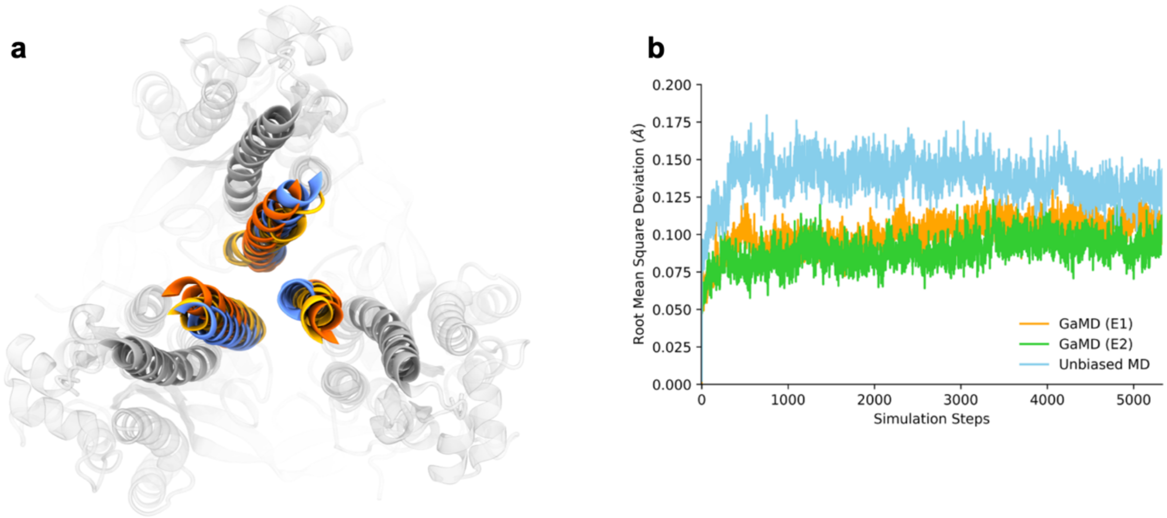


**Supplementary Fig. 1. Conformations from cMD and GaMD simulations. a**, Top-down view of SARS-CoV-2 spike HexaPro-SS-Δstalk in the open prefusion conformation and select conformations from all-atom conventional MD and GaMD simulations. The open crystal structure is shown in gray, with the central helices highlighted as fully opaque. The central helices of alternative simulation methods are shown in color; all-atom conventional MD is shown in blue, and GaMD at the high (E=2) and low (E=1) acceleration level is shown in yellow and orange, respectively. The conformations from all-atom MD and GaMD simulations were selected from the frame with the greatest distance between P987$_{C\alpha}$-P987$_{C\alpha}$ at the top of the central helices, indicating a departure from the closed state en route to a partially open conformation. **b**, Comparative analysis of Root Mean Square Deviation (RMSD) values for three different MD simulation conditions: GaMD (E1), GaMD (E2), and all-atom conventional MD (Unbiased MD). The RMSD values, measured in Angstroms (Å), provide insights into the structural differences in the trajectories over simulation steps.

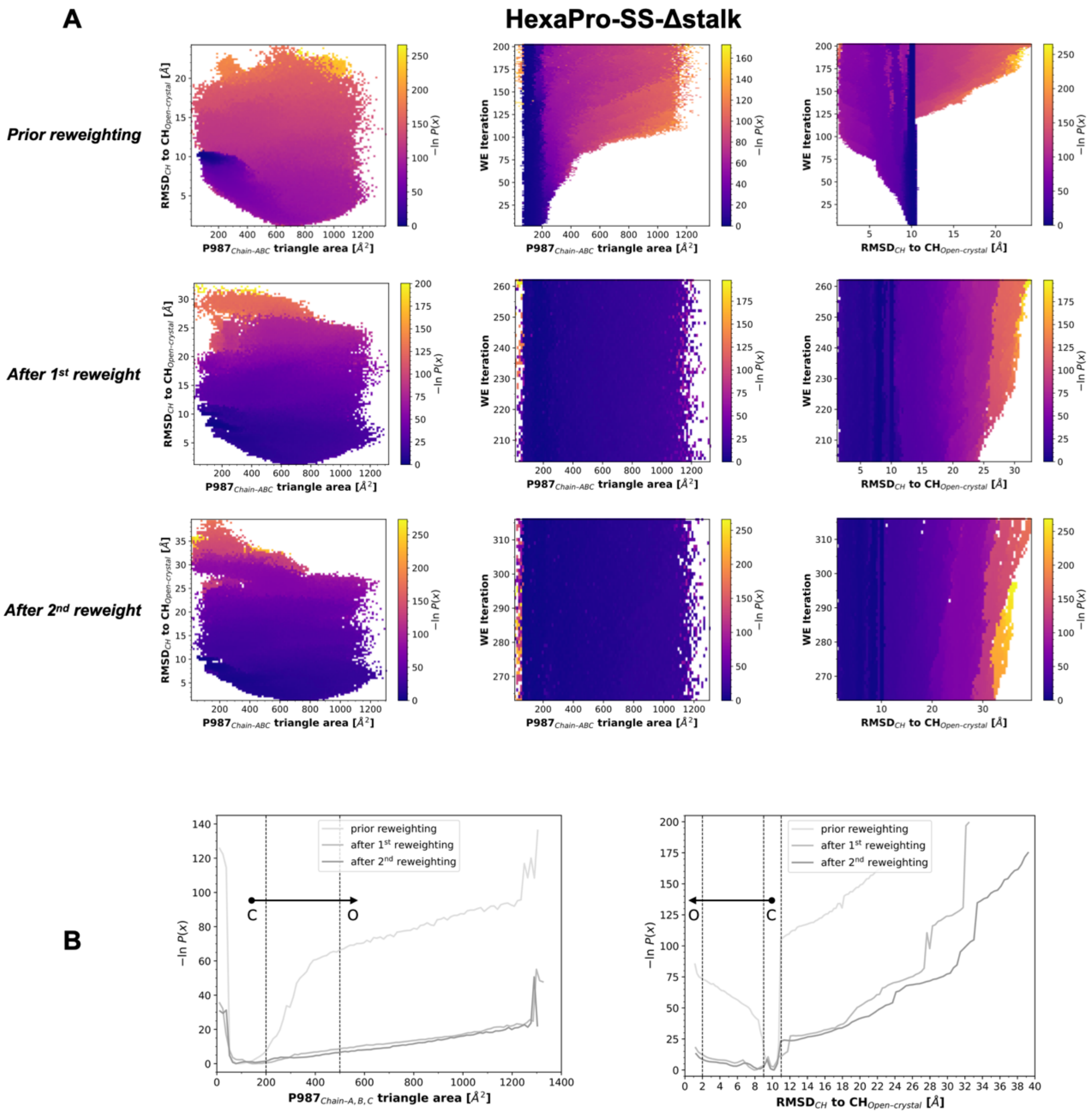


**Supplementary Fig. 2. Probability distribution of progress coordinates in the HexaPro-SS-Δstalk WE simulation. a,** Probability distributions of WE progress coordinates from WE iterations preceding reweighting, in between 1st and 2nd reweighting, and after 2nd reweighting are shown for HexaPro-SS-Δstalk. The probability distribution is plotted on an inverted natural log scale [i.e., -*ln*P(x)], corresponding to the free energy in 1/$k_B$T unit. From left to right, the probability is shown for the two progress coordinates (x-axis: $P987_{C\alpha}$ triangle area; y-axis: $RMSD_{CH}$) as an average, for the $P987_{C\alpha}$ triangle area-only as a time (i.e., WE iterations) evolution, and for the $RMSD_{CH}$-only as time (i.e., WE iterations) evolution, respectively. **b,** 1-D probability distributions for $P987_{C\alpha}$ triangle area (left panel) and $RMSD_{CH}$ (right panel). Different shades of gray are used to plot the average probability calculated from WE iterations preceding reweighting, in between 1st and 2nd reweighting, and after 2nd reweighting, respectively. Dashed lines demarcate closed and open states. An arrow indicates the closed-to-open transition.

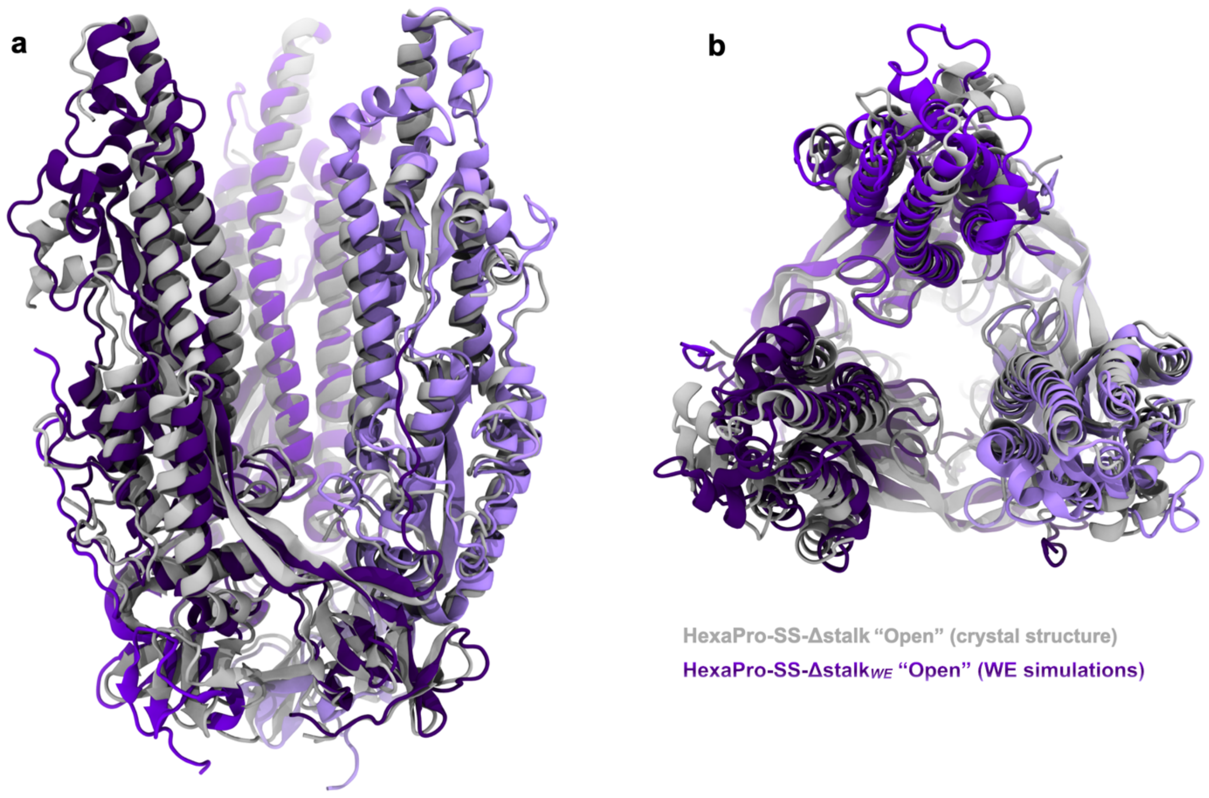


**Supplementary Fig. 3. Overlay of the open S2 trimer crystal structure and open WE model.** The HexaPro-SS-Δstalk S2-only trimer in the open prefusion conformation is depicted with a cartoon representation from side (**a**) and top-down (**b**) perspectives. The open WE model shows protomers highlighted with different shades of purple and the open crystal structure (PDB ID: 8U1G[32]) is shown in gray.

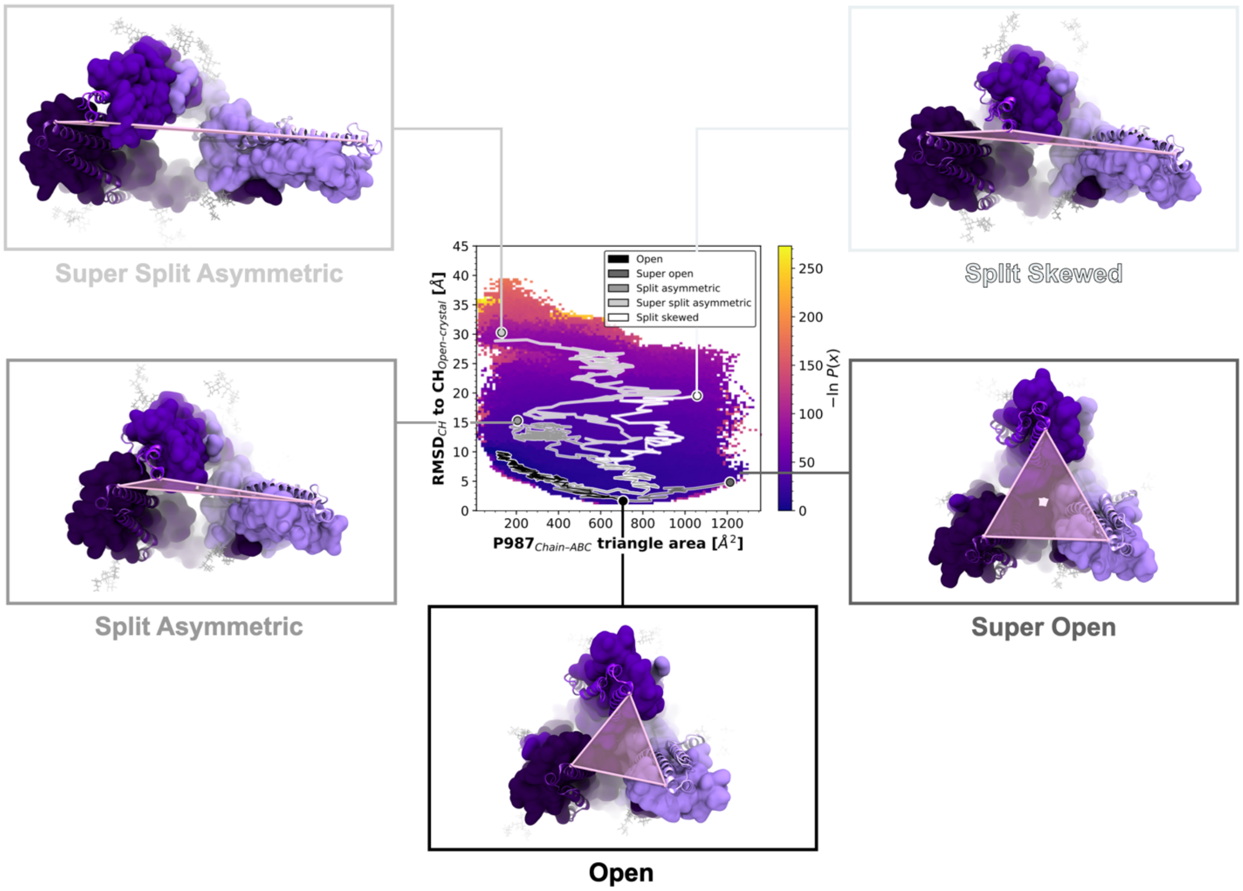


**Supplementary Fig. 4. Additional open conformation sampled during WE simulations.** Beyond the closed-to-open transitions, WE simulations allowed us to sample further open conformations, that we renamed as Super Open, to Split Asymmetric, Split Skewed, and Super Split Asymmetric. Selected pathways leading to each one of these states are indicated with a line (black to white) on top of the 2D probability distribution plot for the two progress coordinates used in the WE simulations of HexPro-SS-Δstalk (x-axis: P987$_{C\alpha}$ triangle area; y-axis: RMSD$_{CH}$). Respective molecular representations for these additional states are reported around the plot. A *top-down* viewpoint is used, with the protomers colored with different shades of purple. We note that, although Split Asymmetric and Super Split Asymmetric states are located on the left-hand side region of the plot, denoted by low P987$_{C\alpha}$ triangle area values, they should instead be located on the right-end side of the probability plot with larger values of P987$_{C\alpha}$ triangle area. This discrepancy occurred because one apex separated significantly from the other two (>100 Å), extending beyond the central box of the simulation. Consequently, it entered the adjacent periodic image, interacting with the other two apexes from the opposite direction. This caused Amber's *cpptraj* program to see the tree apexes almost as in the closed conformation with low values of P987$_{C\alpha}$ triangle area, thus explaining all the conformations residing above the closed state in the probability distribution plot. We note that we tried to prevent this inconvenience by setting an unusually large simulation box (we added a 25 Å layer of water molecules from the solute edge in each direction), but we could not foresee such an exceedingly large opening (> 100 Å P987–P987 distance).

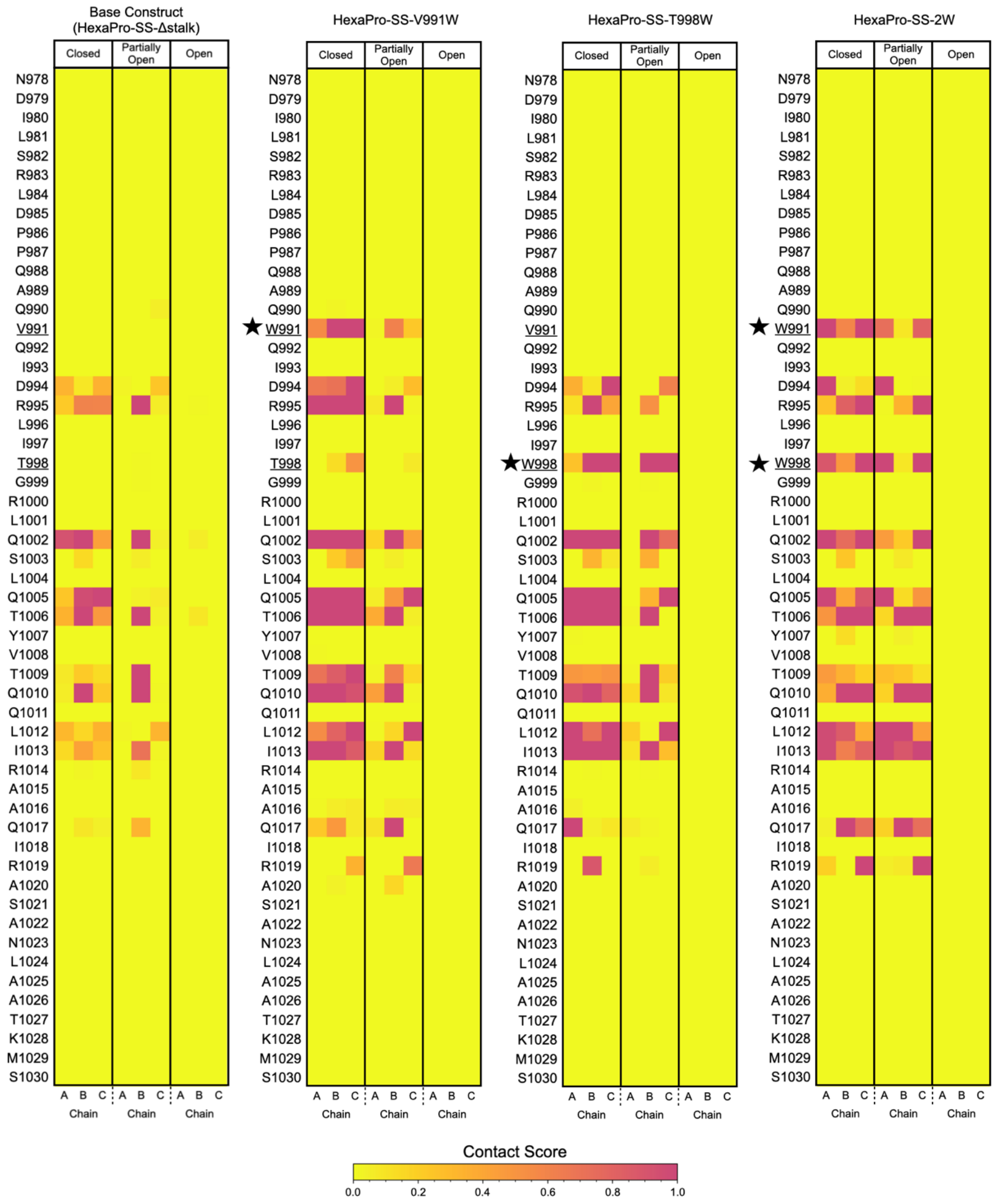


**Supplementary Fig. 5. Contact heatmaps for central helix (CH) residues.** CH contact score heat maps for the HexaPro-SS-Δstalk base construct and HexaPro-SS-Δstalk tryptophan mutants in the closed, partially open, and open conformations. A star indicates a position mutated to tryptophan. Positions 991 and 998 are underlined for each S2 construct. The scale ranges from 0 (weak or transient contacts) to 1 (persistent or extensive contacts), and individual contact scores are normalized to the 95th percentile value of the HexaPro-SS-Δstalk contact scores.

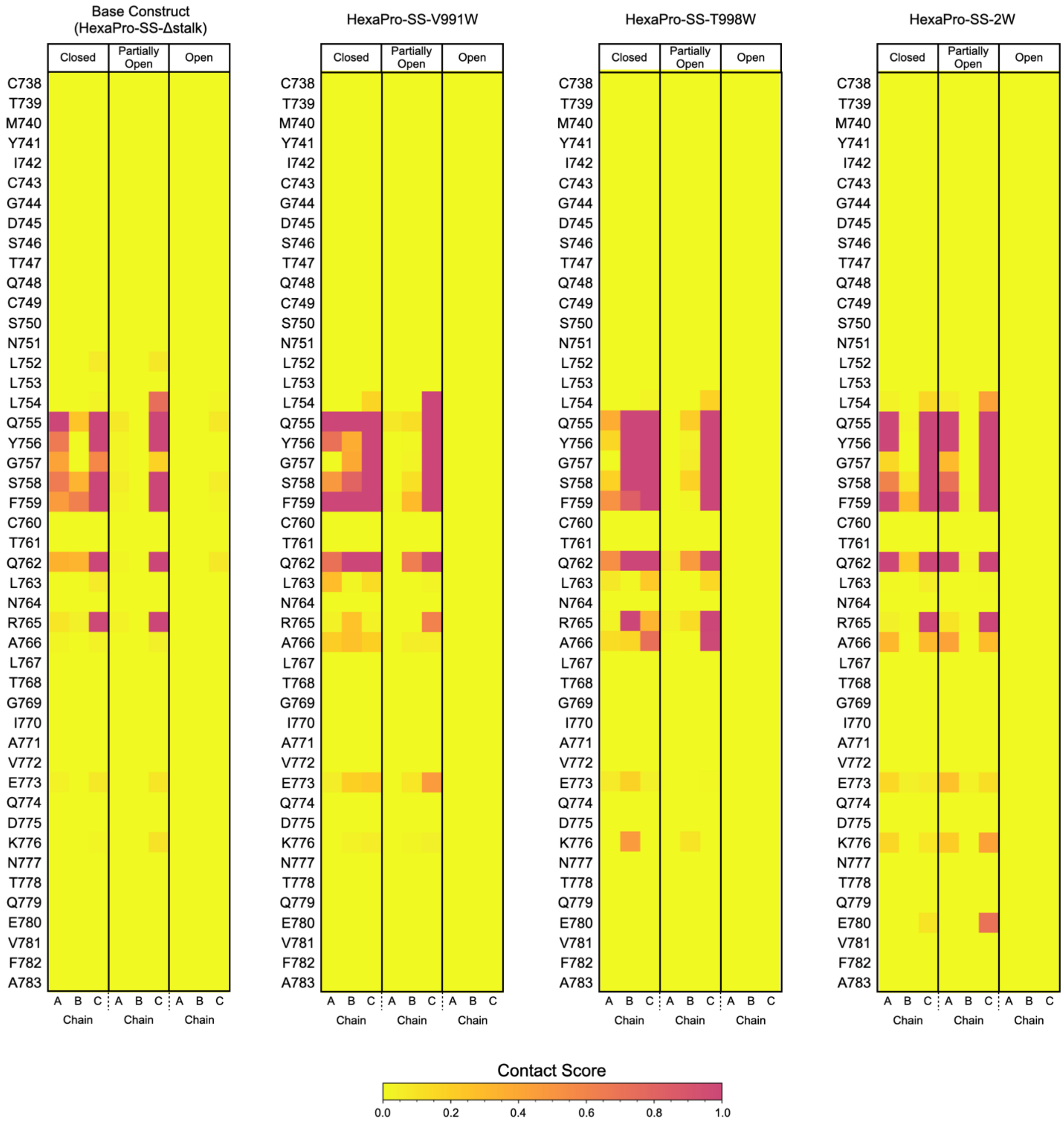


**Supplementary Fig. 6. Contact heatmaps for upper helix (UH) residues.** UH contact score heat maps for the HexaPro-SS-Δstalk base construct and HexaPro-SS-Δstalk tryptophan mutants in the closed, partially open, and open conformations.

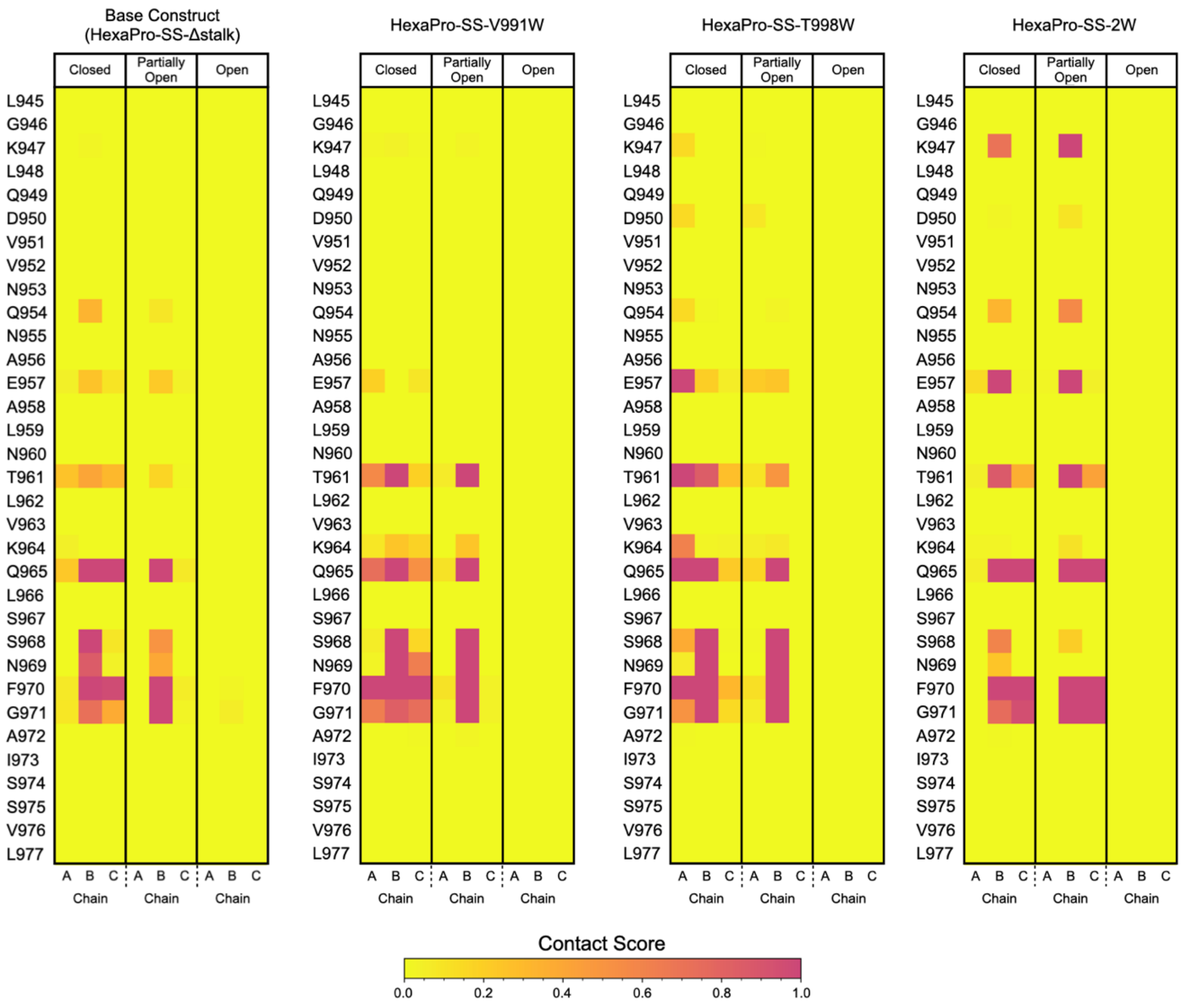


**Supplementary Fig. 7. Contact heatmaps for heptad repeat 1 (HR1) residues.** HR1 contact score heat maps for the HexaPro-SS-Δstalk base construct and HexaPro-SS-Δstalk tryptophan mutants in the closed, partially open, and open conformations.

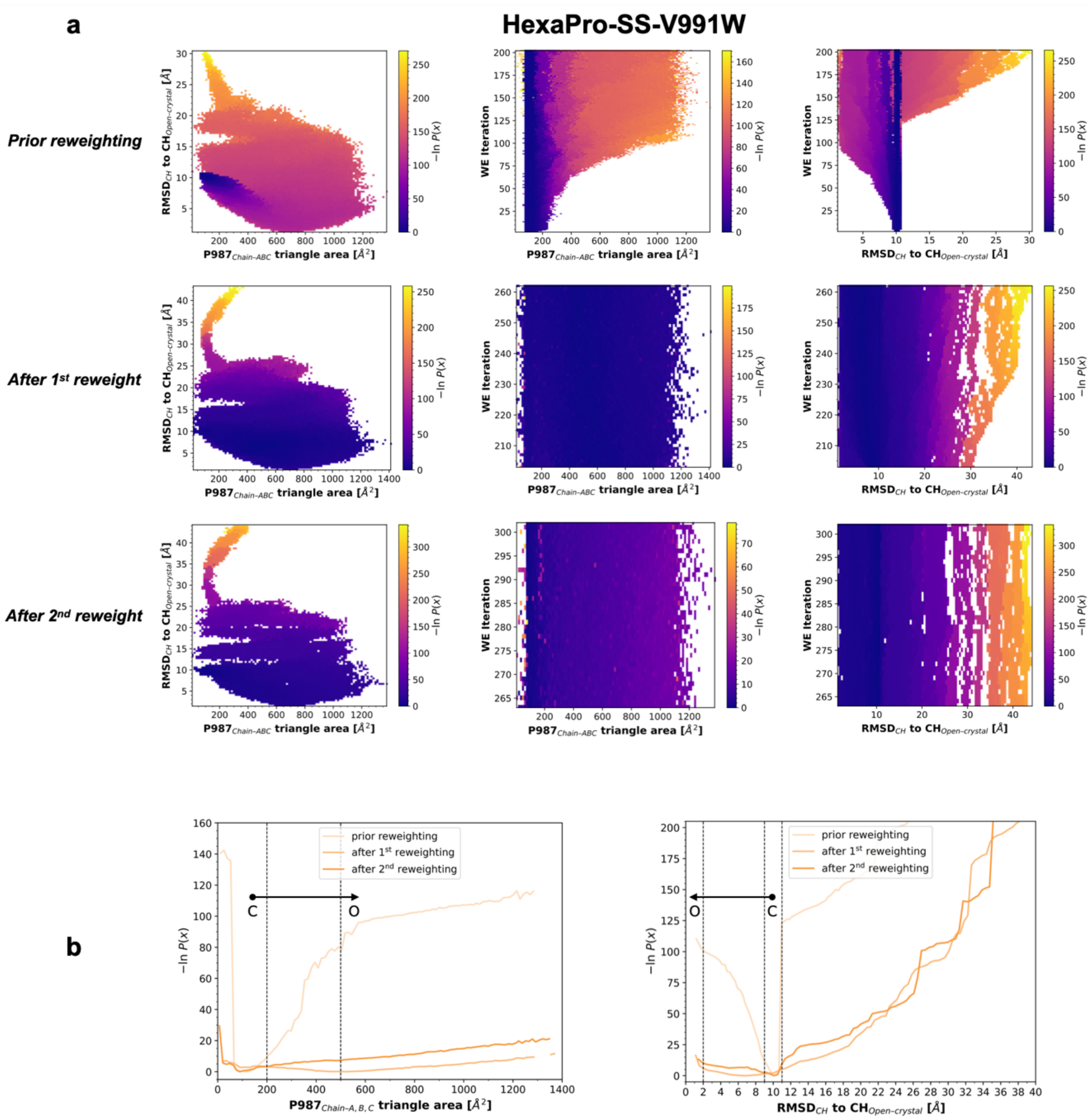


**Supplementary Fig. 8. Probability distribution of progress coordinates in the HexaPro-SS-V991W WE simulations. a,** Probability distributions of WE progress coordinates from WE iterations preceding reweighting, in between 1st and 2nd reweighting, and after 2nd reweighting are shown for HexaPro-SS-V991W. The probability distribution is plotted on an inverted natural log scale [i.e., -*ln*P(x)], corresponding to the free energy in $1/k_BT$ unit. From left to right, the probability is shown for the two progress coordinates (x-axis: $P987_{C\alpha}$ triangle area; y-axis: $RMSD_{CH}$) as an average, for the $P987_{C\alpha}$ triangle area-only as a time (i.e., WE iterations) evolution, and for the $RMSD_{CH}$-only as time (i.e., WE iterations) evolution, respectively. **b,** 1-D probability distributions for $P987_{C\alpha}$ triangle area (left panel) and $RMSD_{CH}$ (right panel). Different shades of orange are used to plot the average probability calculated from WE iterations preceding reweighting, in between 1st and 2nd reweighting, and after 2nd reweighting, respectively. Dashed lines demarcate closed and open states. An arrow indicates the closed-to-open transition.

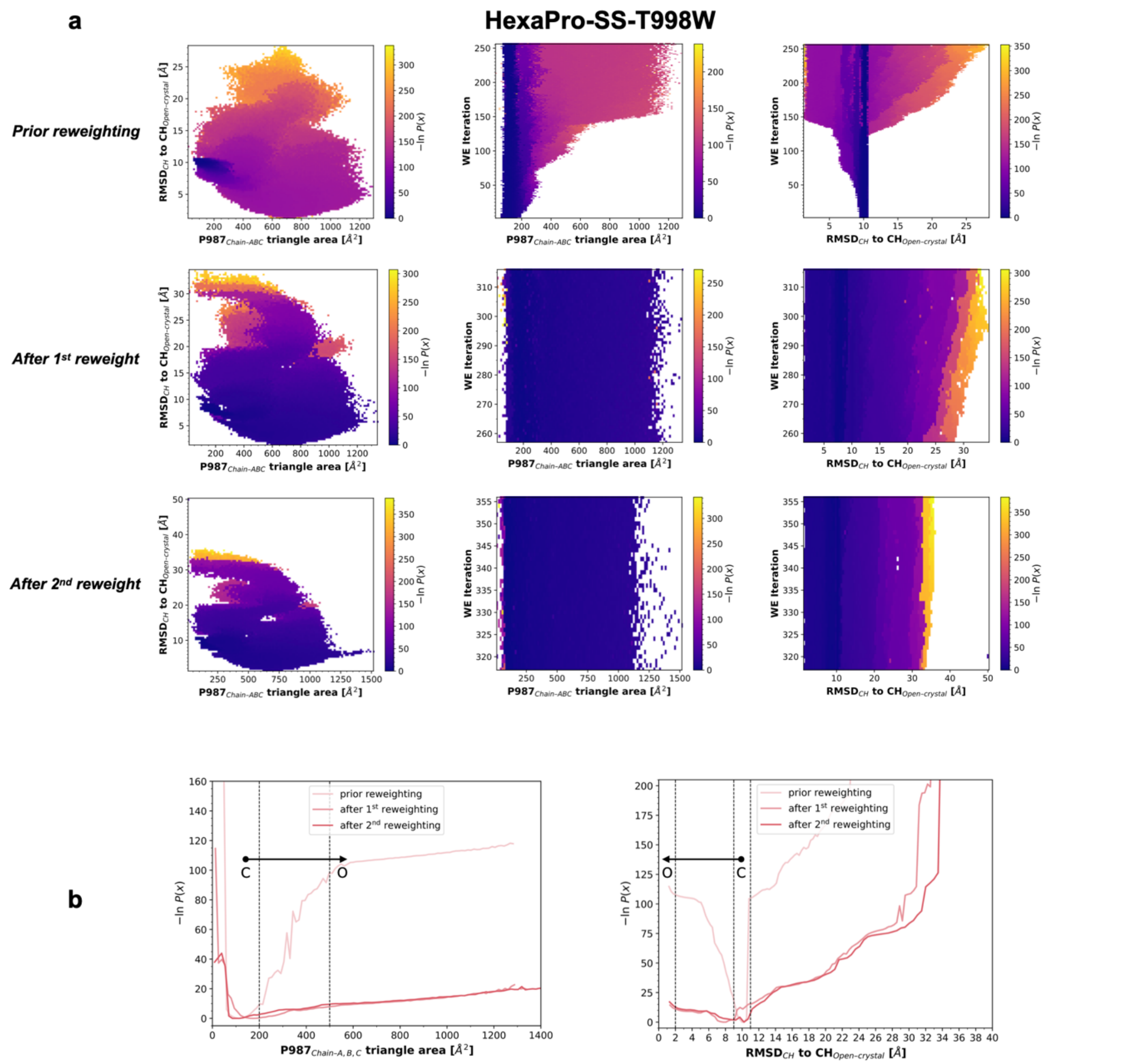


**Supplementary Fig. 9. Probability distribution of progress coordinates in the HexaPro-SS-T998W WE simulations. a,** Probability distributions of WE progress coordinates from WE iterations preceding reweighting, in between 1st and 2nd reweighting, and after 2nd reweighting are shown for HexaPro-SS-T998W. The probability distribution is plotted on an inverted natural log scale [i.e., -*ln*P(x)], corresponding to the free energy in 1/$k_B$T unit. From left to right, the probability is shown for the two progress coordinates (x-axis: P987$_{C\alpha}$ triangle area; y-axis: RMSD$_{CH}$) as an average, for the P987$_{C\alpha}$ triangle area-only as a time (i.e., WE iterations) evolution, and for the RMSD$_{CH}$-only as time (i.e., WE iterations) evolution, respectively. **b,** 1-D probability distributions for P987$_{C\alpha}$ triangle area (left panel) and RMSD$_{CH}$ (right panel). Different shades of red are used to plot the average probability calculated from WE iterations preceding reweighting, in between 1st and 2nd reweighting, and after 2nd reweighting, respectively. Dashed lines demarcate closed and open states. An arrow indicates the closed-to-open transition.

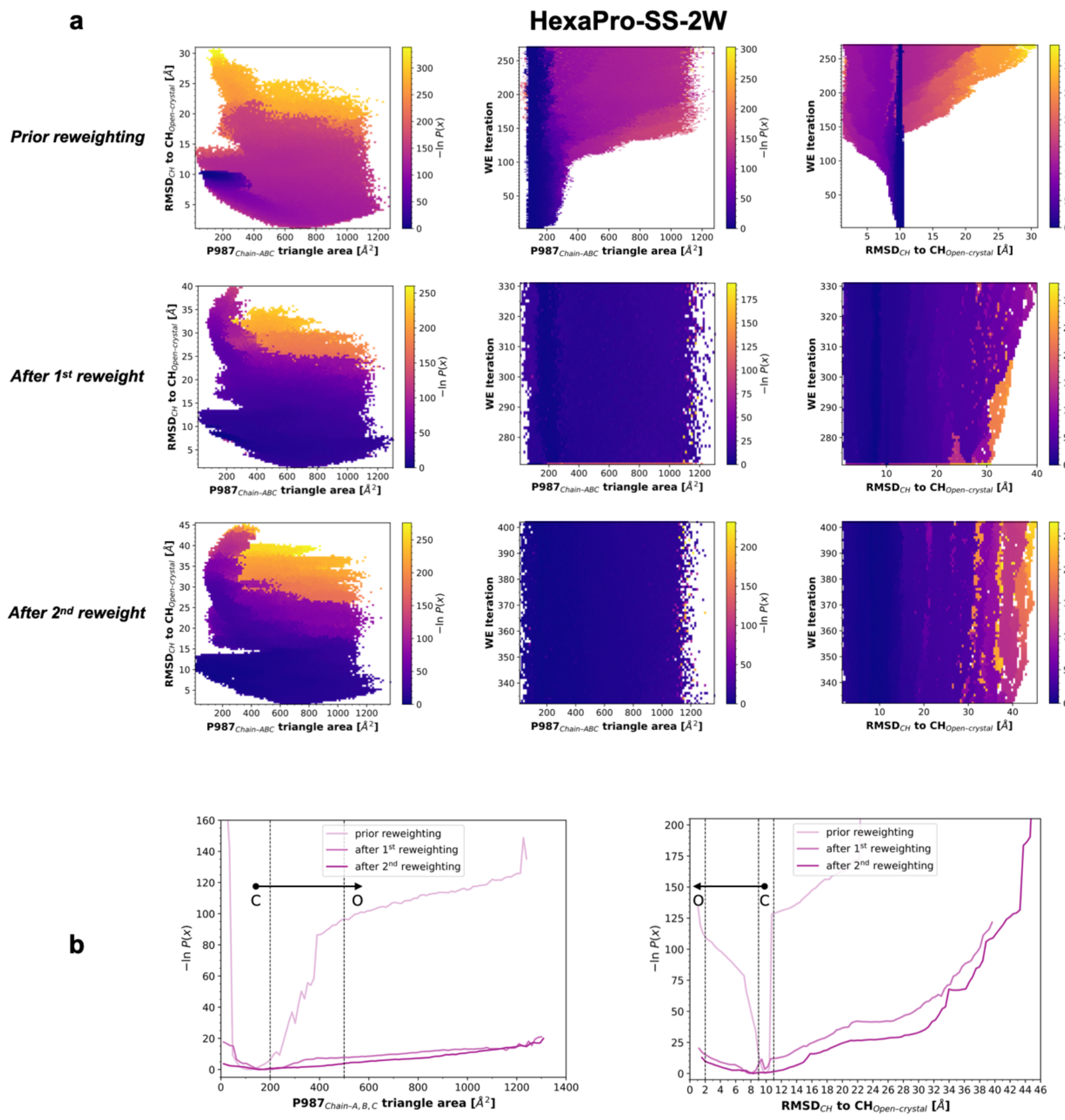


**Supplementary Fig. 10. Probability distribution of progress coordinates in the HexaPro-SS-2W WE simulations. a,** Probability distributions of WE progress coordinates from WE iterations preceding reweighting, in between 1$^{st}$ and 2$^{nd}$ reweighting, and after 2$^{nd}$ reweighting are shown for HexaPro-SS-2W. The probability distribution is plotted on an inverted natural log scale [i.e., -*ln*P(x)], corresponding to the free energy in $1/k_BT$ unit. From left to right, the probability is shown for the two progress coordinates (x-axis: P987$_{C\alpha}$ triangle area; y-axis: RMSD$_{CH}$) as an average, for the P987$_{C\alpha}$ triangle area-only as a time (i.e., WE iterations) evolution, and for the RMSD$_{CH}$-only as time (i.e., WE iterations) evolution, respectively. **b,** 1-D probability distributions for P987$_{C\alpha}$ triangle area (left panel) and RMSD$_{CH}$ (right panel). Different shades of purple are used to plot the average probability calculated from WE iterations preceding reweighting, in between 1$^{st}$ and 2$^{nd}$ reweighting, and after 2$^{nd}$ reweighting, respectively. Dashed lines demarcate closed and open states. An arrow indicates the closed-to-open transition.

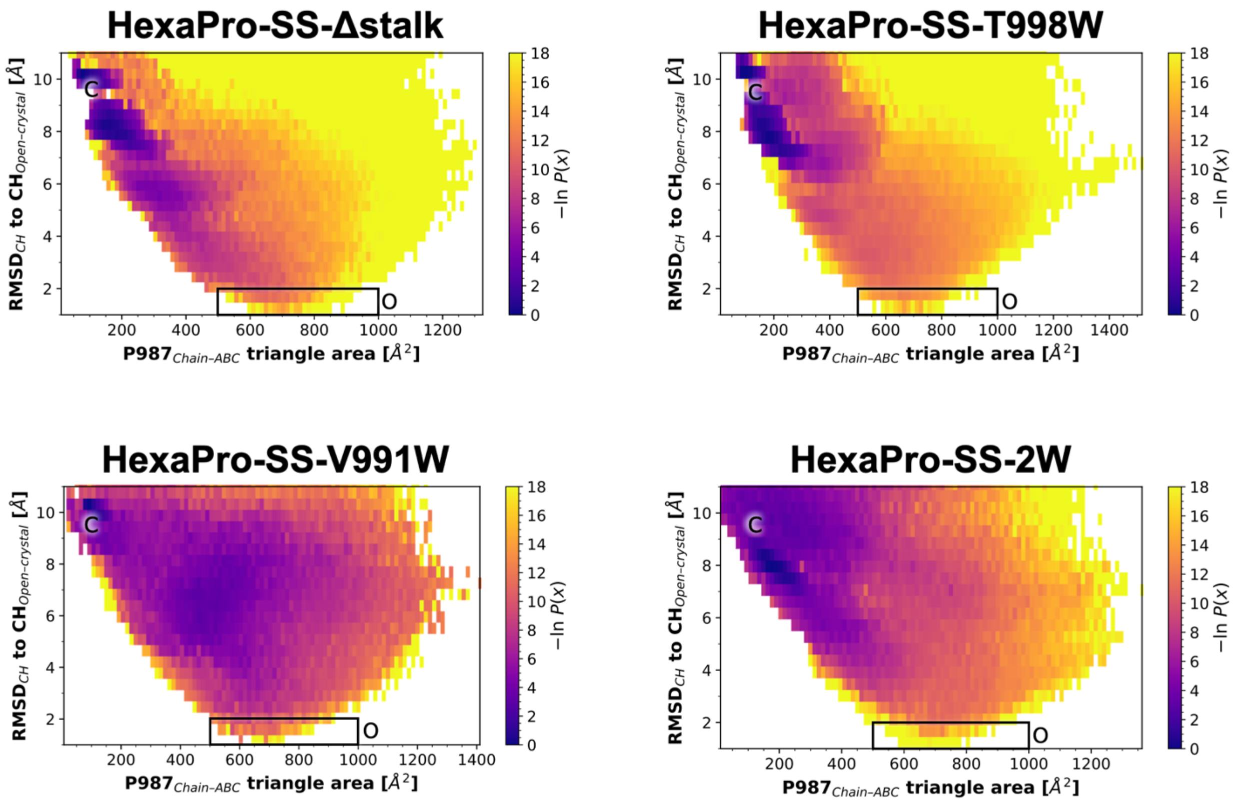


**Supplementary Fig. 11. Probability distribution of progress coordinates within the region-of-interest in the HexaPro-SS-Δstalk, HexaPro-SS-V991W, HexaPro-SS-T998W, and HexaPro-SS-2W WE simulations.** Probability distributions of WE progress coordinates from WE iterations after the 2$^{nd}$ reweighting are shown for HexaPro-SS-Δstalk, HexaPro-SS-V991W, HexaPro-SS-T998W, and HexaPro-SS-2W. The probability distribution is plotted on an inverted natural log scale [i.e., -*ln*P(x)], corresponding to the free energy in 1/$k_B$T unit. The probability is shown for the two progress coordinates (x-axis: P987$_{C\alpha}$ triangle area; y-axis: RMSD$_{CH}$) as an average. The closed state is marked with a "C" and the open state is marked with an "O."

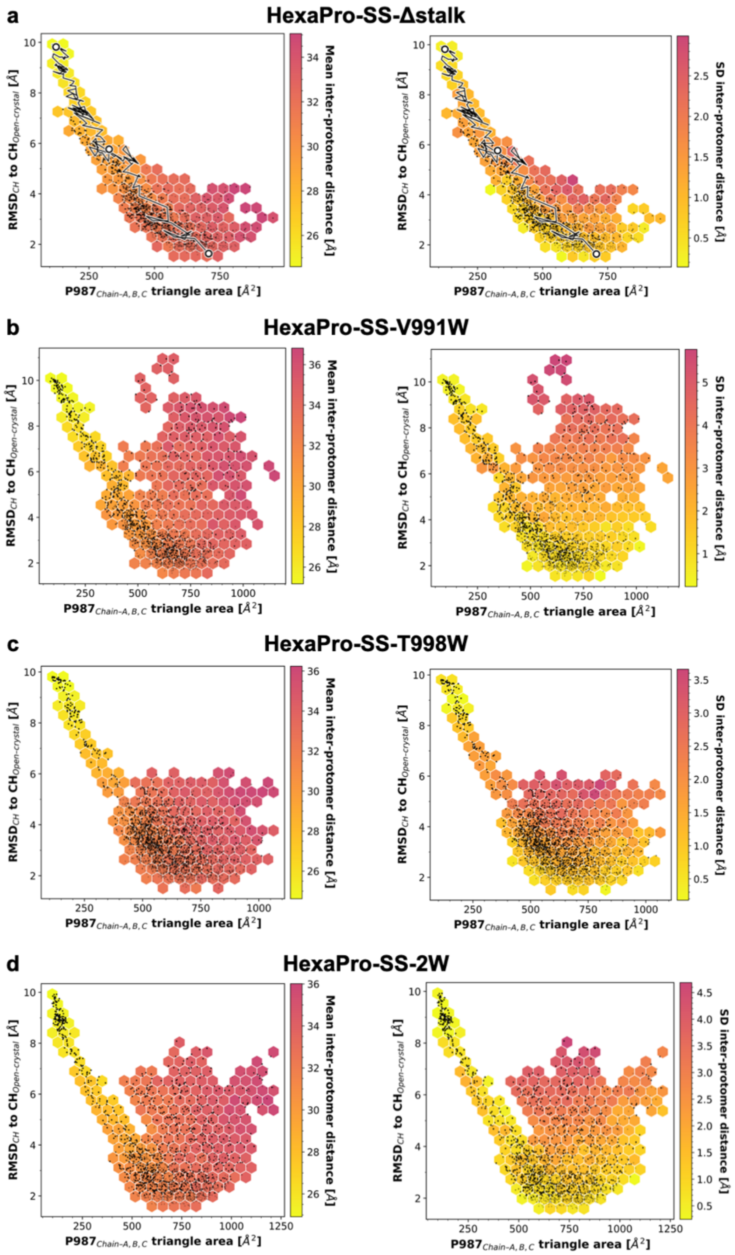


**Supplementary Fig. 12. Conformational space sampled successful pathways obtained from WE simulations.** Distribution of the conformations sampled in the opening pathways obtained from the WE simulation of (**a**) HexaPro-SS-Δstalk (**b**) HexaPro-SS-V991W, (**c**) HexaPro-SS-T998W, (**d**) HexaPro-SS-2W. Each black point represents a conformation sampled along an opening pathway. The RMSD of the CH to $CH_{Open\text{-}crystal}$ (y-axis) is plotted against the area of the triangle formed by the $P987_{C\alpha}$ at the CH apex (x-axis). In the left-side panels, the hexagonal bin color is scaled to the mean interprotomer distance of the black data points within the respective bin, whereas on the right-side panels, the hexagon bin color is scaled according to the standard deviation of the black data points within the respective bin. For (**a**) A trace of one of the successful pathways is shown as a black line with white points corresponding to the closed, partially open, and open conformations.

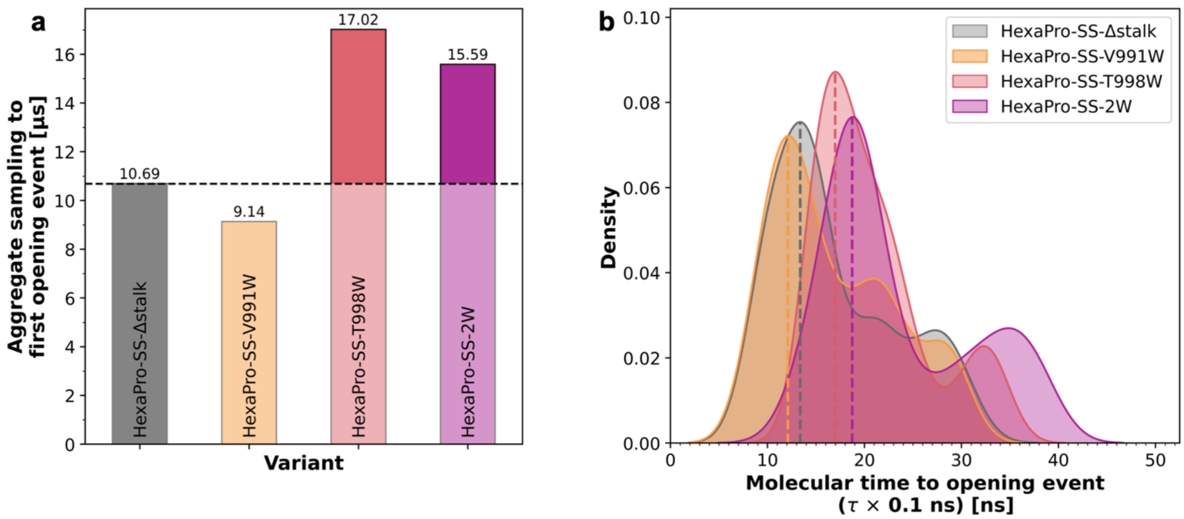


**Supplementary Fig. 13. Aggregate molecular time to first opening and distribution of molecular time to opening. a,** Aggregate molecular time to the first opening event observed in the WE simulations of HexaPro-SS-Δstalk, HexaPro-SS-V991W, HexaPro-SS-T998W, and HexaPro-SS-2W. The number in μs is reported on top of each respective bar. **b,** Distribution of molecular time to opening for all the successful pathways observed in the WE simulations of HexaPro-SS-Δstalk, HexaPro-SS-V991W, HexaPro-SS-T998W, and HexaPro-SS-2W, respectively. The distributions are shown as kernel densities and do not take into account the weights of successful pathways. Differently from the opening event duration (Fig. S12), the molecular time to opening event (indicated in units of ns) takes into account the time spent in the closed state. A dashed line indicates the peak of each distribution.

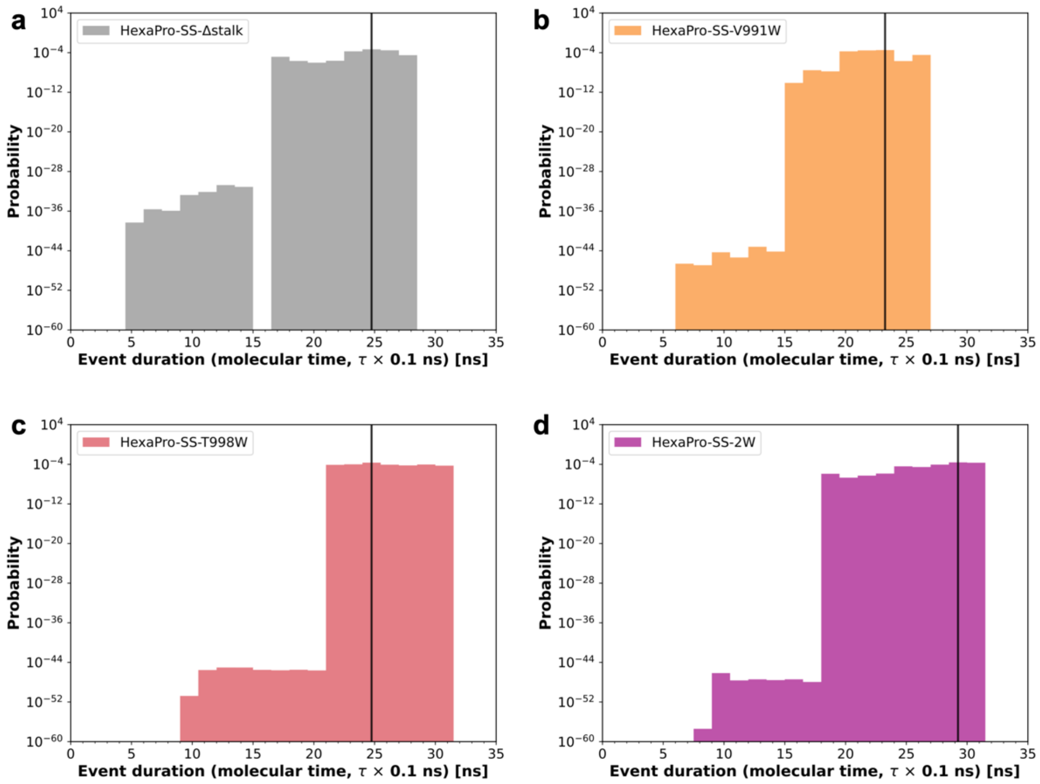


**Supplementary Fig. 14. Probability of S2 opening event duration.** Probability of S2 trimer opening event duration as calculated from the opening pathways observed in the WE simulations of (**a**) HexaPro-SS-Δstalk, (**b**) HexaPro-SS-V991W, (**c**) HexaPro-SS-T998W, and (**d**) HexaPro-SS-2W. The probability considers the weight of the successful pathways calculated from WE simulations. Differently from the molecular time to opening event (Fig. S11), the duration of the opening event does not take into account the time spent in the closed state, but rather the effective molecular time required by the system to reach the open state after leaving the closed state. A solid black line indicates the most likely event duration.

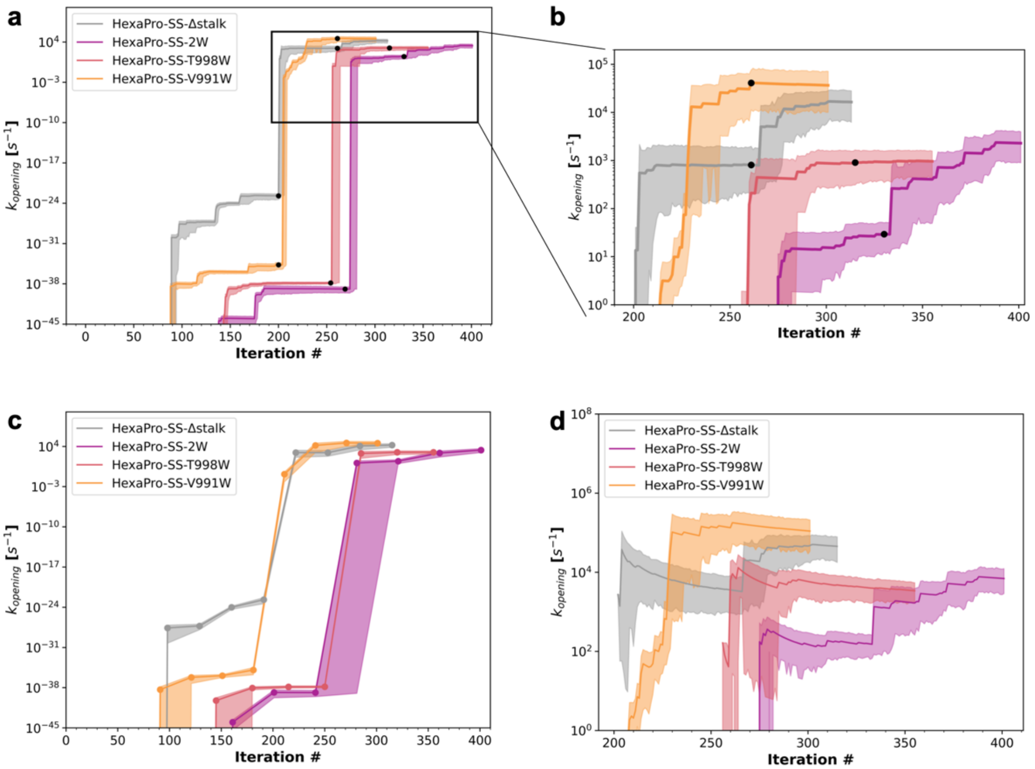


**Supplementary Fig. 15. Rate constants of S2 trimer closed-to-open (*opening*) transition. a-d**, Time (i.e., iteration #) evolution of rate constant estimates ($k_{opening}$) in units of $s^{-1}$ of S2 trimer closed-to-open (opening) transition as calculated from WE simulations of HexaPro-SS-Δstalk, HexaPro-SS-V991W, HexaPro-SS-T998W, and HexaPro-SS-2W. The uncertainty in the rate constants is illustrated with a semi-transparent shaded region that extends from the lower to the upper bounds of the confidence interval (95%) for the observable. Solid black circles are marked at the iterations where reweighting took place. **a**, Time (i.e., iteration #) evolution of rate constant estimates ($k_{opening}$) in units of $s^{-1}$ of S2 trimer closed-to-open (opening) assessed as a rolling average calculated across all WE iterations over 1-iteration windows with 95% confidence interval. **b,** Magnified view of the rate constant evolution after iteration #200 as calculated in panel **a**. **c,** Time (i.e., iteration #) evolution of rate constant estimates ($k_{opening}$) in units of $s^{-1}$ of S2 trimer closed-to-open (opening) assessed as a rolling average calculated across all WE iterations over iteration windows of width corresponding to the 10% of the total number of iterations for the respective WE simulation with 95% confidence interval. **d,** Time (i.e., iteration #) evolution of rate constant estimates ($k_{opening}$) in units of $s^{-1}$ of S2 trimer closed-to-open (opening) assessed as a rolling average calculated for all WE iterations following first reweighting over 1-iteration windows with 95% confidence interval.

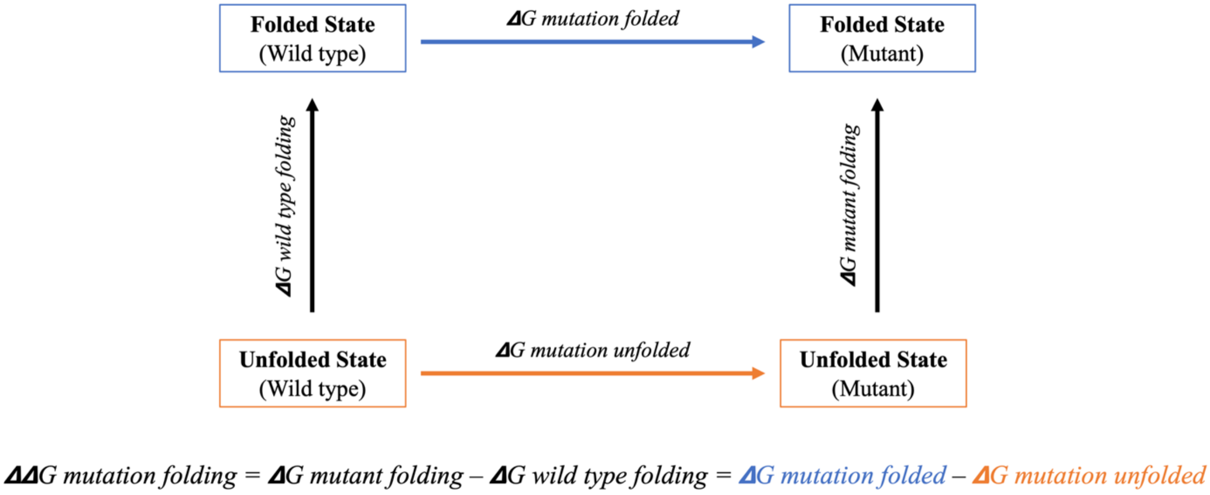


**Supplementary Fig. 16. Schematic representation of a thermodynamic cycle used to calculate changes in protein folding free energy upon mutation (*ΔΔG mutation folding*).** The left column depicts the folding process of a wild-type protein, with the associated folding free energy as *ΔG wild type folding*. The same folding process is presented on the right column, but in the case of a mutated protein, with the associated folding free energy *ΔG mutant folding*. The reaction shown in the top row corresponds to the alchemical transformation of the folded wild type protein into the folded mutant, resulting in the free energy difference between the two forms as *ΔG mutation folded*. The process depicted in the bottom row represents the same alchemical transformation (wild type into mutant), but in the case of the protein in the unfolded state, with the associated free energy difference *ΔG mutation unfolded*. While computing the free energy differences for the vertical processes is computationally demanding, those for the horizontal transformations are more accessible. Therefore, *ΔΔG mutation folding* can be calculated by taking the difference between *ΔG mutation folded* and *ΔG mutation unfolded*.

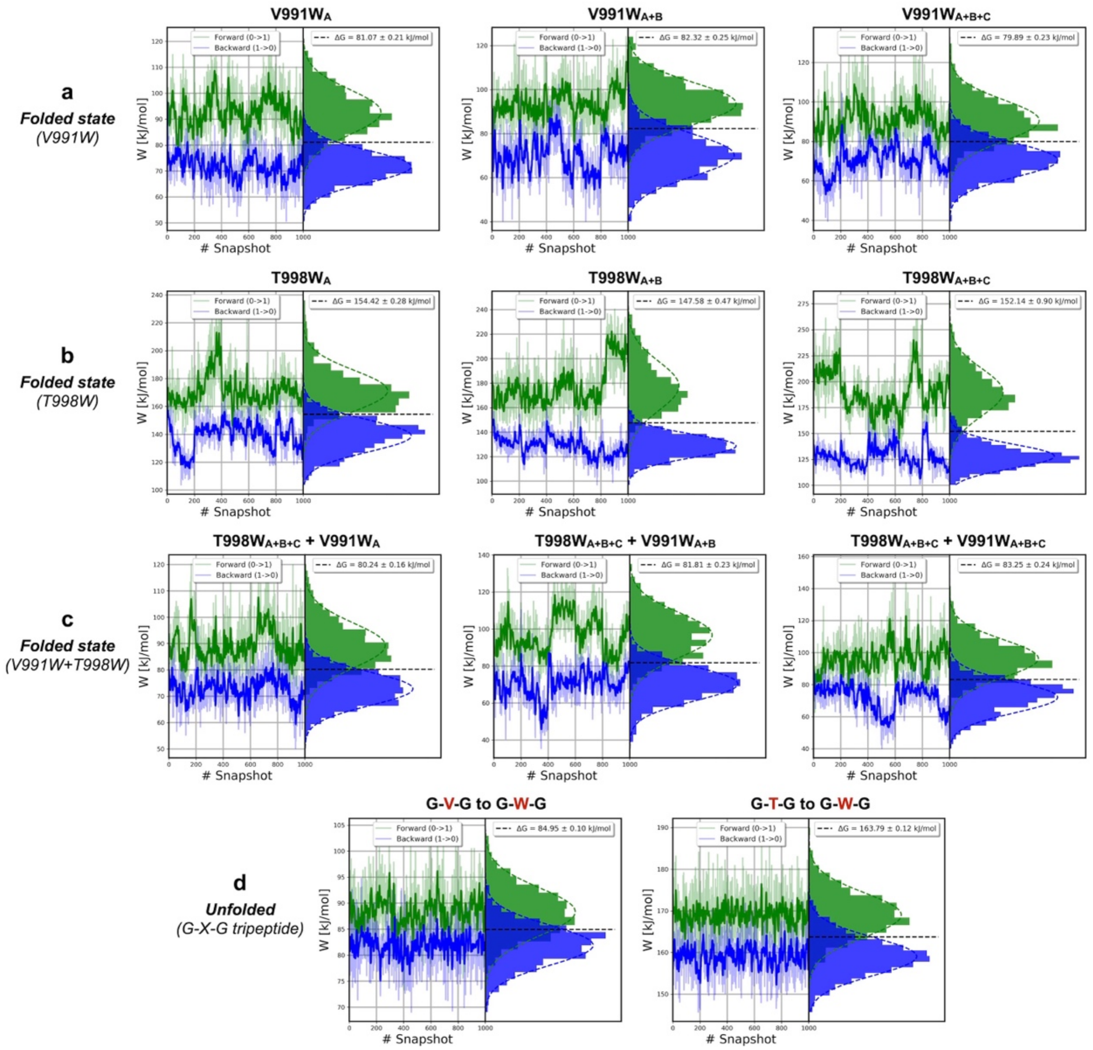


**Supplementary Fig. 17. Alchemical non-equilibrium free energy calculations.** The output of PMX[47] *analyze* script for non-equilibrium MD simulations performed for (**a**) HexaPro-SS-V991W (folded state), (**b**) HexaPro-SS-T998W (folded state), and (**c**) HexaPro-SS-2W (folded state), (**d**) G-V-G / G-T-G tripeptides (unfolded state). In (**a**) and (**b**), the left, central, and right plots account for the V991W or T998W mutation introduced in Chains A, A+B, A+B+C, respectively. In (**c**), the left, central, and right plots account for the V991W mutation introduced in Chains A, A+B, A+B+C, respectively, on top of a base construct already incorporating T998W in all three Chains. In each plot, the left side displays the work values corresponding to both the forward and reverse transitions for each starting structure. 1000 non-equilibrium MD simulations were run in the forward direction (wild type-to-mutant, highlighted in green) and 1000 non-equilibrium MD simulations in the reverse direction (mutant-to-wild type, highlighted in blue) for both the folded and the unfolded state. During these calculations, one amino acid is gradually morphed into the mutated amino acid (forward) or vice-versa (reverse). On the right side of the plots, histograms illustrate the distributions of these work values, and their overlap allows the determination of the free energy difference.

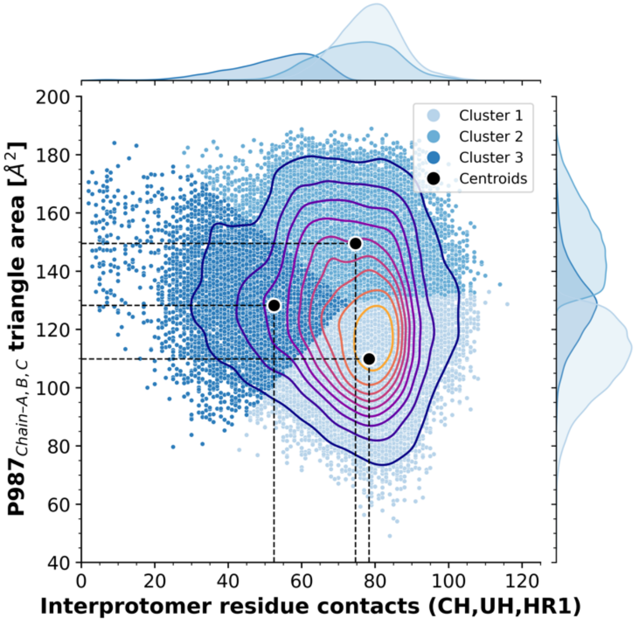


**Supplementary Fig. 18. Cluster analysis of the closed conformations extracted from WE simulation of HexaPro-SS-2W.** All the closed conformations sampled during the WE simulations of HexaPro-SS-2W were clustered using *k-means* clustering based on the number of interprotomer contacts made by CH, UH, and HR1 helices (x-axis) and the $P987_{TRIANGLE\text{-}AREA}$ observables. The three identified clusters are highlighted with different shades of blue. Centroids for each cluster are marked with solid black circles. Dashed lines indicate the values of the observables corresponding to the identified centroids. The population density is emphasized through contour lines illustrated with a plasma color palette, with the area encircled by the yellow contour representing the highest population density. Individual population densities for each cluster are shown on the top x-axis for the number of interprotomer contacts and on the right y-axis for the $P987_{TRIANGLE\text{-}AREA}$.

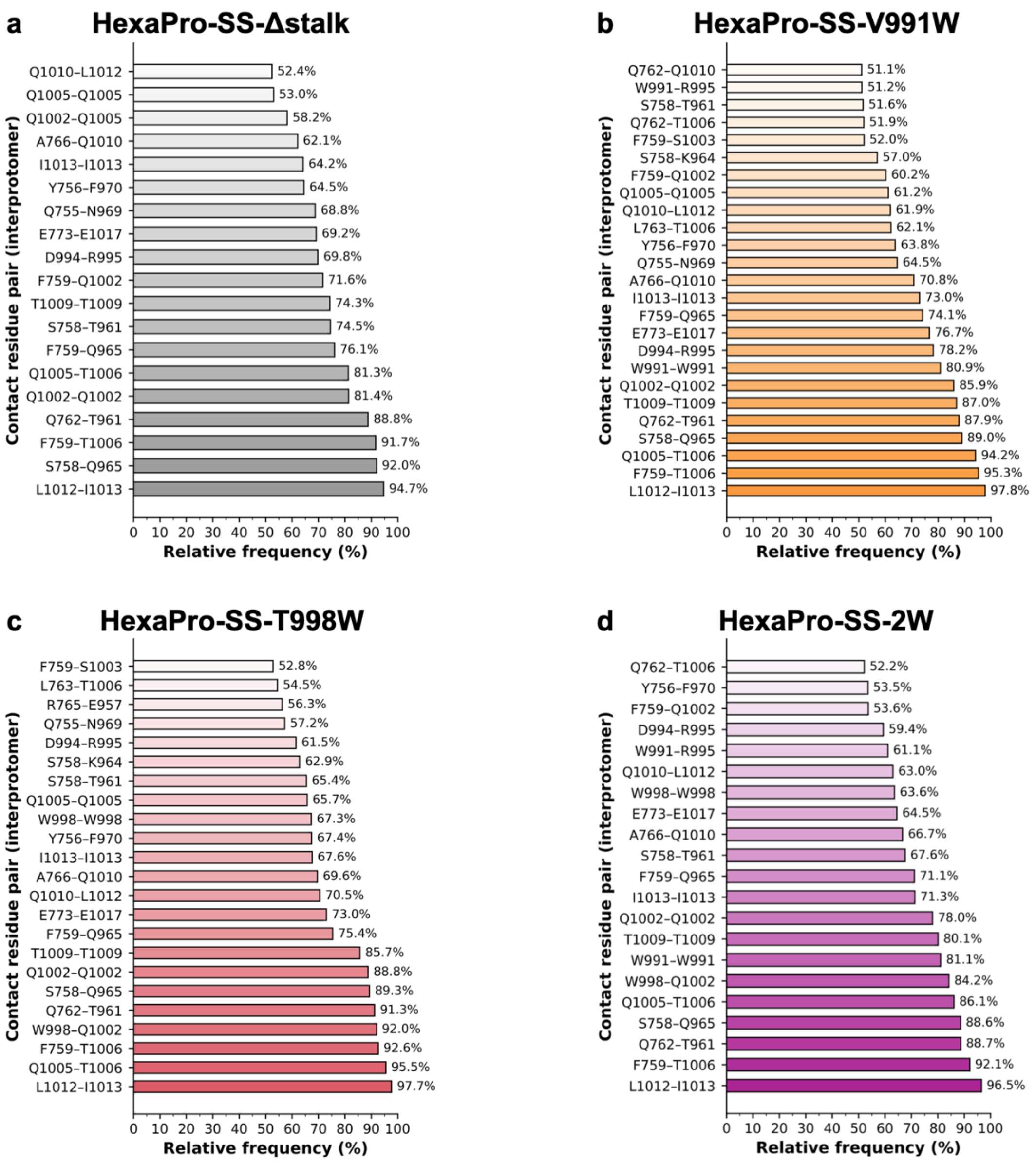


**Supplementary Fig. 19. Contact analysis of the closed conformations extracted from WE simulations.** The bar plots show the relative frequency of occurrence (x-axis, % to the total number of frames × $3_{\text{protomers}}$) of residue pairs within CH, UH, HR1 involved in interprotomer contacts (y-axis) across all the closed conformations extracted from the WE simulations of (**a**) HexaPro-SS-Δstalk, (**b**) HexaPro-SS-V991W, (**c**) HexaPro-SS-T998W, (**d**) and HexaPro-SS-2W. Only contacts with a relative frequency larger than 50% are shown. Contacts are estimated based on a distance cutoff of 4.5 Å between heavy atoms only.

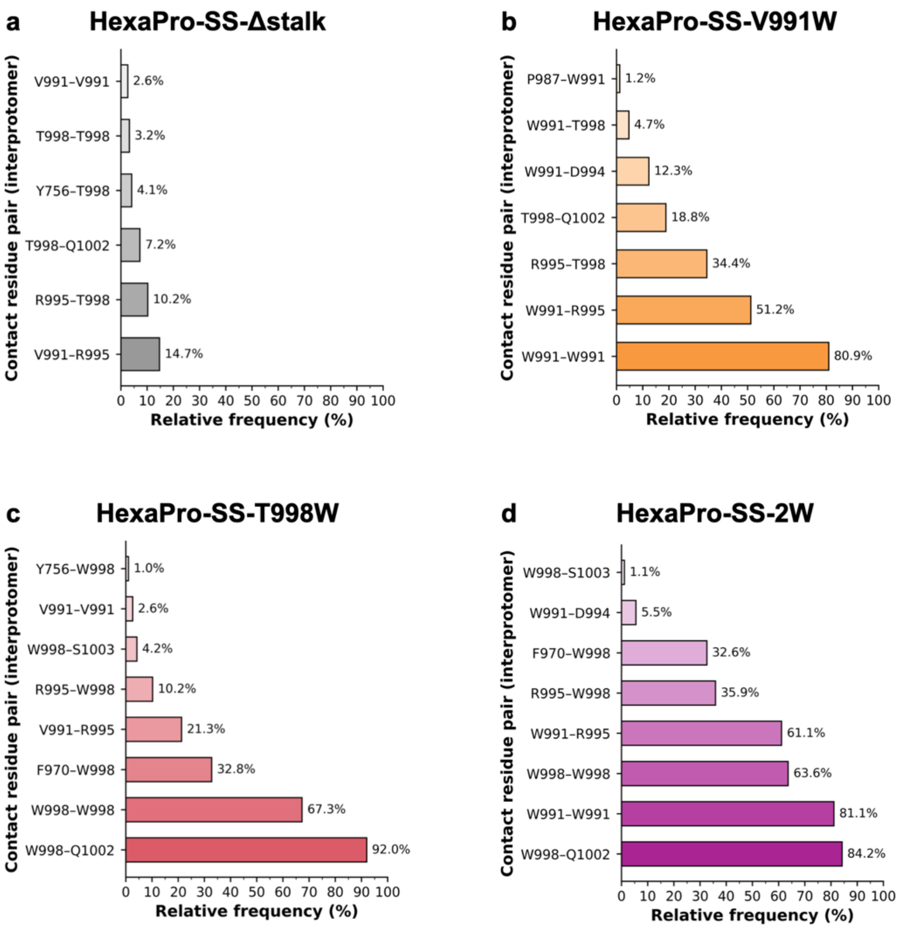


**Supplementary Fig. 20. Contact analysis of the closed conformations extracted from WE simulations (residues 991 and 998 only).** The bar plots show the relative frequency of occurrence (x-axis, % to the total number of frames × $3_{protomers}$) of residue pairs that involve residues at position 991 and 998 forming interprotomer contacts (y-axis) across all the closed conformations extracted from the WE simulations of (**a**) HexaPro-SS-Δstalk, (**b**) HexaPro-SS-V991W, (**c**) HexaPro-SS-T998W, (**d**) and HexaPro-SS-2W. Only the contacts with a relative frequency larger than 1% are shown. Contacts are estimated based on a distance cutoff of 4.5 Å between heavy atoms only.

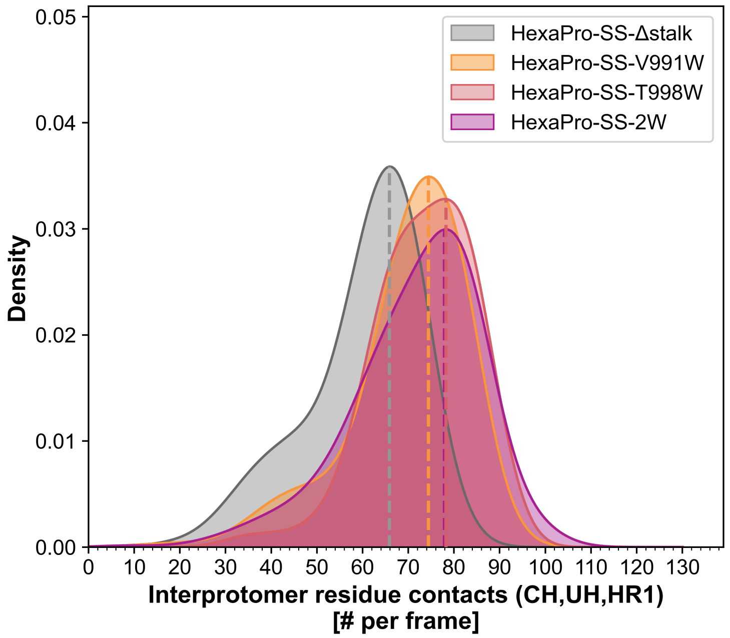


**Supplementary Fig. 21. Number of interprotomer residue contacts in the closed conformations from WE simulations.** The distribution of the number of interprotomer contacts per frame involving residues within CH, UH, and HR1 helices is plotted for HexaPro-SS-Δstalk (gray), HexaPro-SS-V991W (orange), HexaPro-SS-T998W (red), and HexaPro-SS-2W (purple) using kernel density curves. The contacts are assessed exclusively from the closed conformations extracted from their respective WE simulation and are calculated based on a distance cutoff of 4.5 Å considering only the heavy atoms. Dashed lines are drawn from the respective distribution peak.

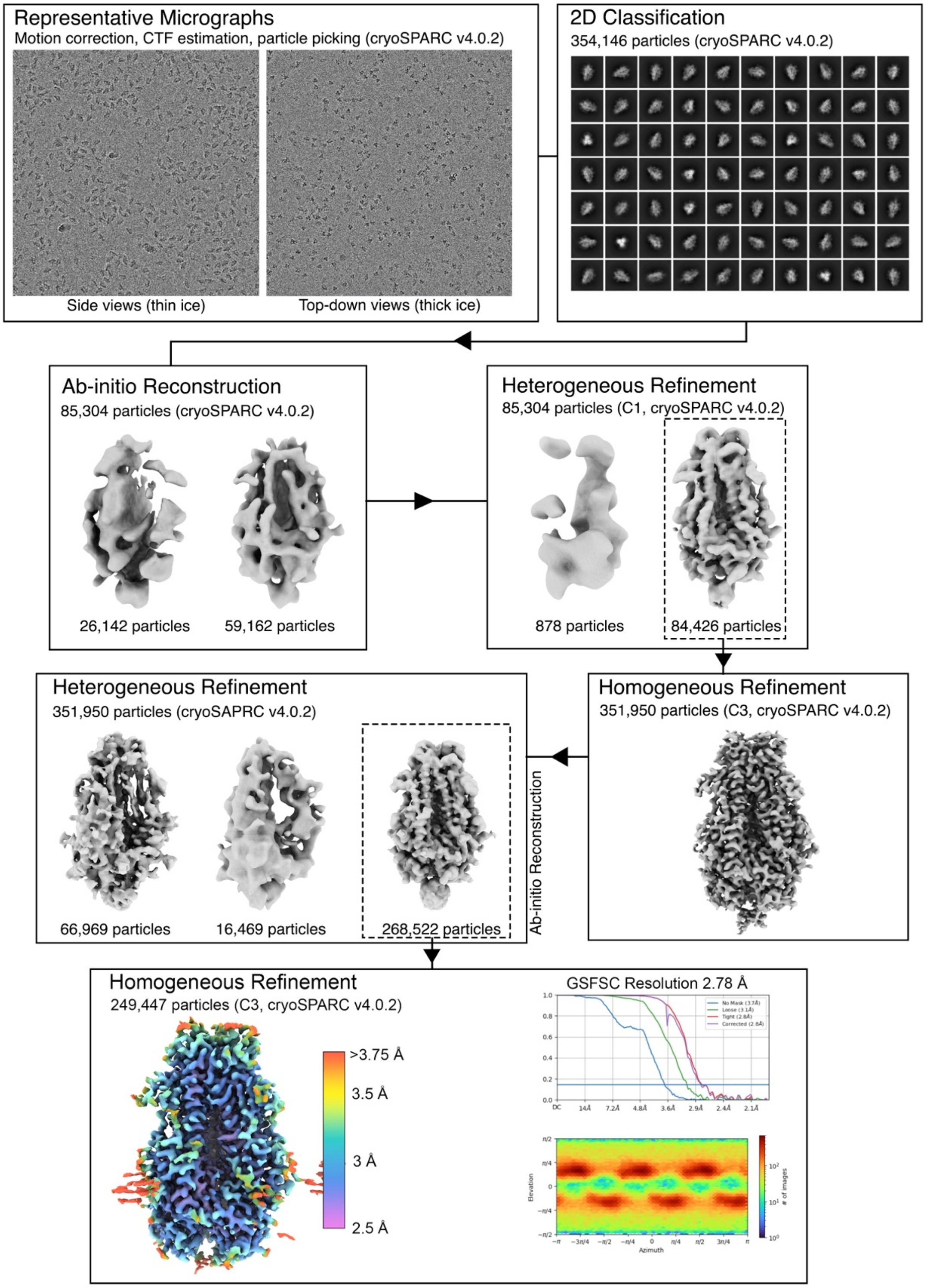


**Supplementary Fig. 22. Cryo-EM data processing workflow for HexaPro-SS-2W.** The following steps were performed in cryoSPARC[59] v4.0.2 to process the cryo-EM dataset: motion correction, CTF estimation, 2D classification, ab-initio reconstruction, heterogeneous refinement, and homogeneous refinement (C3 symmetry).

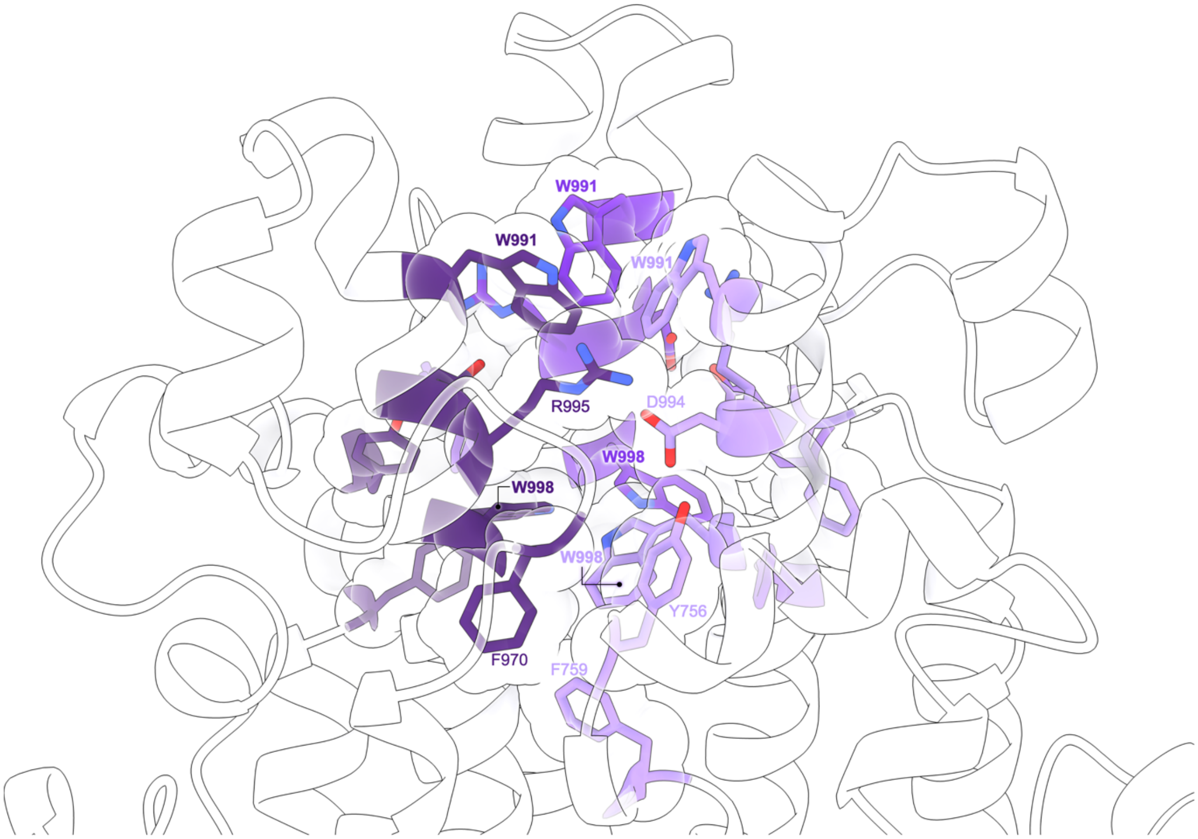


**Supplementary Fig. 23. Hydrophobic packing in the cryo-EM structure of HexaPro-SS-2W.** Sidechains of residues contributing to forming hydrophobic packing and D994–R995 salt bridge at the S2 trimer apex are illustrated as sticks colored in varying tones of purple according to the respective chain (A being the lightest, C being the darkest). For these residues, a transparent sphere representation is also overlaid. The rest of the protein is shown as transparent cartoons.

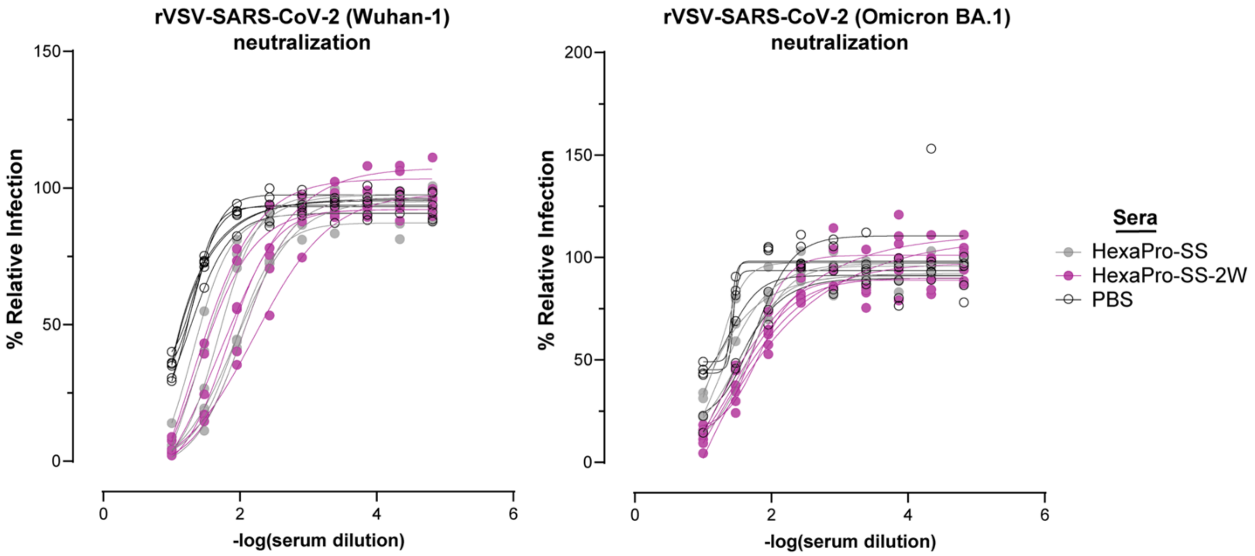


**Supplementary Fig. 24. Individual replicates of rVSV-SARS-CoV-2 neutralization assays.** Female C57BL/6J mice (n = 6) were immunized with HexaPro-SS, HexaPro-SS-2W (V991W+T998W), or PBS. Serial 3-fold dilutions were made for each serum sample and assayed for neutralizing ability against rVSV-SARS-CoV-2 (Wuhan-1), on the left, or (Omicron BA.1), on the right, in singlicate. One mouse immunized with HexaPro-SS needed to be euthanized for humane reasons before sufficient blood could be collected. As a result, the corresponding neutralization assay against rVSV-SARS-CoV-2 (Omicron BA.1) for that immunization group only had n = 5. Infectivity curves were fitted with nonlinear regression (variable slope; four parameters; GraphPad Prism 9.3.1). Area under the curve values were calculated using GraphPad Prism 9.3.1.

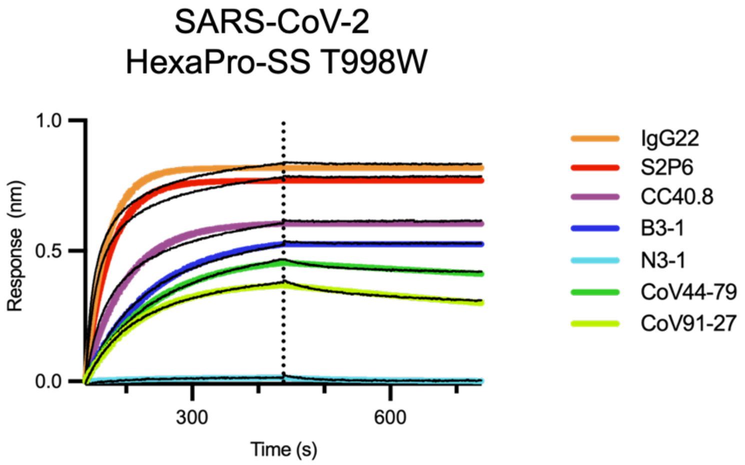


**Supplementary Fig. 25.** Biolayer interferometry sensorgrams showing binding of previously reported neutralizing or non-neutralizing antibodies to epitopes on the S2 subunit in the SARS-CoV-2 HexaPro-SS-T998W construct (including stalk region). Colored lines indicate 1:1 binding fit, black lines indicate reference subtracted response.

### 3. Supplementary Tables 1 – 4

**Supplementary Table 1.** Summary of the systems that were simulated in this work. List of abbreviations: conventional MD simulations (cMD), Gaussian accelerated MD simulations (GaMD), weighted ensemble MD simulations (WE), alchemical non-equilibrium free energy calculations (FEC).

| Method | System | Box dimensions (Å x Å x Å) | Total atoms (# Atoms) | Salt Concentration (M) | Aggregate Sampling Time (µs) |
|---|---|---|---|---|---|
| cMD | HexaPro-SS | 134 Å x 140 Å x 153 Å | 274,082 | 0.15 | 2.63 |
| GaMD | HexaPro-SS | 134 Å x 140 Å x 153 Å | 274,082 | 0.15 | 3.53 |
| WE | HexaPro-SS | 138 Å x 150 Å x 168 Å | 328,200 | 0.15 | 45.79 |
| | HexaPro-SS-V991W | 138 Å x 150 Å x 168 Å | 328,328 | 0.15 | 35.72 |
| | HexaPro-SS-T998W | 138 Å x 150 Å x 168 Å | 328,271 | 0.15 | 47.66 |
| | HexaPro-SS-2W | 138 Å x 150 Å x 168 Å | 328,370 | 0.15 | 56.62 |
| FEC | HexaPro-SS-V991W | 150 Å x 150 Å x 150 Å | 308,022 | 0.15 | 0.60 |
| | HexaPro-SS-T998W | 150 Å x 150 Å x 150 Å | 308,007 | 0.15 | 0.60 |
| | HexaPro-SS-2W | 150 Å x 150 Å x 150 Å | 308,022 | 0.15 | 0.60 |
| | Unfolded (G-V-G) | 50 Å x 50 Å x 50 Å | 5,936 | 0.15 | 0.60 |
| | Unfolded (G-T-G) | 50 Å x 50 Å x 50 Å | 5,946 | 0.15 | 0.60 |
| TOTAL | - | - | - | - | 194.95 |

**Supplementary Table 2.** GaMD boost potential statistics for simulations performed at low (E=1) and high acceleration (E=2).

| Boost | Boost Parameter | Low Acceleration | High Acceleration |
|---|---|---|---|
| Total Potential Boost | $\sigma_{0P}$ (kcal/mol) | 175.0 | 350.0 |
| | $\sigma_{\Delta VP}$ (kcal/mol) | 1734.44 | 2,344.54 |
| | $\Delta V_{Pavg}$ (kcal/mol) | -866676 | 862577 |
| | $k_{0P}$ | 5.33 x $10^{-5}$ | 6.15 x $10^{-5}$ |
| Dihedral Potential Boost | $\sigma_{0D}$ (kcal/mol) | 15.0 | 35.0 |
| | $\sigma_{\Delta VD}$ (kcal/mol) | 83.24 | 124.36 |
| | $\Delta V_{Davg}$ (kcal/mol) | 15991.6 | 16194.5 |
| | $k_{0D}$ | 9.86 x $10^{-4}$ | 9.68 x $10^{-4}$ |

**Supplementary Table 3**. Summary of observable quantities calculated from WE and non-equilibrium alchemical free energy simulations. The table includes for each simulated system, from left to right: rate constants in $s^{-1}$ ($k_{opening}$) extracted from all WE iterations, from WE iterations following reweighting, Mean First Passage Time (ms) calculated from the rate constants using the Hill relation, *ΔΔG mutation folding* (kcal/mol) calculated from alchemical non-equilibrium free energy calculations with respect to the HexaPro-SS-Δstalk (see Fig. S14), total aggregate sampling (μs) from WE simulations, total wall time for WE simulations (days) corresponding to the actual number of days required to complete the WE simulations, total number of opening pathways observed during WE simulations, mean duration of an opening event ± standard error of the mean.

| Variant | $k_{opening}$ [full simulation] ($s^{-1}$) | $k_{opening}$ [post reweighting] ($s^{-1}$) | MFPT (ms) | ΔΔG *mutation folding* (kcal/mol) | Total sampling (μs) | Total wall time (days) | Opening pathways (number) | Duration of opening event (mean ± s.e.m.) |
|---|---|---|---|---|---|---|---|---|
| **HexaPro-SS-Δstalk** | $(1.6 \pm 0.6) \times 10^4$ | $(4.5 \pm 1.5) \times 10^4$ | 0.022 | – | 45.7864 | 45.36 | 180 | 137.1 ± 4.8 ns |
| **HexaPro-SS-V991W** | $(3.7 \pm 1.5) \times 10^4$ | $(1.1 \pm 0.5) \times 10^5$ | 0.009 | -2.76 ± 0.11 | 35.7192 | 45.67 | 149 | 139.5 ± 5.0 ns |
| **HexaPro-SS-T998W** | $(9.6 \pm 3.0) \times 10^2$ | $(3.4 \pm 0.1) \times 10^3$ | 0.294 | -8.91 ± 0.26 | 47.6576 | 58.38 | 187 | 168.7 ± 4.1 ns |
| **HexaPro-SS-2W** | $(2.3 \pm 0.8) \times 10^3$ | $(7.0 \pm 2.3) \times 10^3$ | 0.143 | -12.18 ± 0.34 | 56.6160 | 54.49 | 195 | 154.9 ± 5.0 ns |
| **TOTAL** | – | | | | **185.7792** | **203.9** | **711** | – |

**Supplementary Table 4. EM data collection for HexaPro-SS-2W-Δstalk**

| **EM data collection** | |
|---|---|
| Microscope | FEI Glacios |
| Voltage (kV) | 200 |
| Detector | Falcon 4 |
| Magnification (nominal) | 150,000 |
| Pixel size (Å/pix) | 0.94 |
| Exposure rate ($e^-$/pix/sec) | 3.79 |
| Exposure ($e^-$/$Å^2$) | 50 |
| Defocus range (μm) | 1.5-2.5 |
| Tilt angle (˚) | 0 |
| Micrographs collected | 1,764 |
| Micrographs used | 1,622 |
| Particles extracted (total) | 744,038 |
| Automation software | SerialEM |
| Sample | HexaPro-SS-2W-Δstalk |

| | |
|---|---|
| Particles | 249,447 |
| Symmetry | C3 |
| Map sharpening B-factor | -107.3 |
| Unmasked resolution at 0.5 FSC (Å) | 3.2 |
| Masked resolution at 0.5 FSC (Å) | 3.0 |
| Unmasked resolution at 0.143 FSC (Å) | 2.8 |
| Masked resolution at 0.143 FSC (Å) | 2.8 |

| **Model refinement and validation statistics** | |
|---|---|
| Refinement package | ChimeraX ISOLDE, Phenix |
| Refinement tool | Phenix real-space refinement |
| Refinement strategies | min global, local_grid_search, adp, occupancy, reference model restraints, ss restraints, rotamer restraints, Ramachandran restraints, NCS constraints |
| Composition | |
| Amino acids | 1311 |
| RMSD bonds (Å) | 0.003 |
| RMSD angles (º) | 0.49 |
| Average B-factors | |
| Amino acids | 53.3 |
| Ramachandran | |
| Favored (%) | 97.7 |
| Allowed (%) | 2.3 |
| Outliers (%) | 0 |
| Rotamer outliers (%) | 0 |
| Clash score | 2.86 |
| C-beta outliers (%) | 0 |
| CaBLAM outliers (%) | 2.08 |
| CC (mask) | 0.77 |
| MolProbity score | 1.14 |
| EMRinger score | 4.19 |

### 4. Captions for Supplementary Movies 1,2

**Supplementary Movie 1. Opening of HexaPro-SS-Δstalk S2 trimer.** The movie shows the closed-to-open transition of HexaPro-SS-Δstalk as obtained from the respective WE MD simulation. Top-down (left) and side (right) views are provided. The S2 trimer is illustrated with a surface representation, where the three protomers are colored with varying shades of purples. The CHs and HR1 helices are depicted as cartoons. Glycans are shown with gray sticks. The opening pathway shown in the movie corresponds to the pathway highlighted in Figure 2.

**Supplementary Movie 2. Stabilization of the S2 trimer's closed prefusion conformation in HexaPro-SS-2W.** The movie highlights the stabilizing role played by V991W and T998W introduced in HexaPro-SS-2W. An unsuccessful pathway, i.e., a WE MD trajectory where only closed conformations were sampled, extracted from the respective WE simulation, is also shown. The S2 trimer is illustrated with cartoons colored in varying tones of purple according to the respective chain (A being the lightest, C being the darkest). Side chains of residues contributing to forming hydrophobic packing and D994–R995 salt bridge at the S2 trimer apex are illustrated as sticks. Glycans are shown with gray sticks. Solid lines connect residues that form hydrophobic (π–π) interactions, whereas dashed lines indicate electrostatic (cation–π and salt bridge) interactions.

### 5. Supplementary References

1. Hsieh, C.-L. *et al.* Structure-based design of prefusion-stabilized SARS-CoV-2 spikes. *Science* **369**, 1501–1505 (2020).

2. Bangaru, S. *et al.* Structural analysis of full-length SARS-CoV-2 spike protein from an advanced vaccine candidate. *Science* **370**, 1089–1094 (2020).

3. Humphrey, W., Dalke, A. & Schulten, K. VMD: Visual molecular dynamics. *J. Mol. Graph.* **14**, 33–38 (1996).

4. Ao, J. *et al.* psfgen User's Guide. http://www.ks.uiuc.edu/Research/vmd/plugins/psfgen/ug.pdf (2020).

5. Casalino, L. *et al.* Beyond shielding: The roles of glycans in the SARS-CoV-2 spike protein. *ACS Cent. Sci.* **6**, 1722–1734 (2020).

6. Watanabe, Y., Allen, J. D., Wrapp, D., McLellan, J. S. & Crispin, M. Site-specific glycan analysis of the SARS-CoV-2 spike. *Science* **369**, 330–333 (2020).

7. Shajahan, A., Supekar, N. T., Gleinich, A. S. & Azadi, P. Deducing the N-and O-glycosylation profile of the spike protein of novel coronavirus SARS-CoV-2. *Glycobiology* **30**, 981–988 (2020).

8. Olsson, M. H. M., Søndergaard, C. R., Rostkowski, M. & Jensen, J. H. PROPKA3: Consistent Treatment of Internal and Surface Residues in Empirical p K a Predictions. *J. Chem. Theory Comput.* **7**, 525–537 (2011).

9. Huang, J. *et al.* CHARMM36m: an improved force field for folded and intrinsically disordered proteins. *Nat. Methods* **14**, 71–73 (2017).

10. Huang, J. & MacKerell Jr, A. D. CHARMM36 all-atom additive protein force field: Validation based on comparison to NMR data. *J. Comput. Chem.* **34**, 2135–2145 (2013).

11. Guvench, O., Hatcher, E., Venable, R. M., Pastor, R. W. & MacKerell, A. D. J.

CHARMM Additive All-Atom Force Field for Glycosidic Linkages between Hexopyranoses. *J. Chem. Theory Comput.* **5**, 2353–2370 (2009).

12. Jorgensen, W. L., Chandrasekhar, J., Madura, J. D., Impey, R. W. & Klein, M. L. Comparison of simple potential functions for simulating liquid water. *J. Chem. Phys.* **79**, 1087 (1983).

13. Phillips, J. C. *et al.* Scalable molecular dynamics on CPU and GPU architectures withNAMD. *J. Chem. Phys.* **153**, 44130 (2020).

14. Ermak, D. L. & McCammon, J. A. Brownian dynamics with hydrodynamic interactions. *J. Chem. Phys.* **69**, 1352–1360 (1978).

15. Martyna, G. J., Tobias, D. J. & Klein, M. L. Constant pressure molecular dynamics algorithms. *J. Chem. Phys.* **101**, 4177–4189 (1994).

16. Ryckaert, J.-P., Ciccotti, G. & Berendsen, H. J. C. Numerical integration of the cartesian equations of motion of a system with constraints: molecular dynamics of n-alkanes. *J. Comput. Phys.* **23**, 327–341 (1977).

17. Darden, T., York, D. & Pedersen, L. Particle mesh Ewald: An N·log(N) method for Ewald sums in large systems. *J. Chem. Phys.* **98**, 10089–10092 (1993).

18. Miao, Y., Feher, V. A. & McCammon, J. A. Gaussian Accelerated Molecular Dynamics: Unconstrained Enhanced Sampling and Free Energy Calculation. *J. Chem. Theory Comput.* **11**, 3584–3595 (2015).

19. Wang, J. *et al.* Gaussian accelerated molecular dynamics: Principles and applications. *WIREs Comput. Mol. Sci.* **11**, e1521 (2021).

20. Zuckerman, D. M. & Chong, L. T. Weighted Ensemble Simulation: Review of Methodology, Applications, and Software. *Annu. Rev. Biophys.* **46**, 43–57 (2017).

21. Huber, G. A. & Kim, S. Weighted-ensemble Brownian dynamics simulations for protein association reactions. *Biophys. J.* **70**, 97–110 (1996).

22. Sztain, T. *et al.* A glycan gate controls opening of the SARS-CoV-2 spike protein. *Nat. Chem. 2021 1310* **13**, 963–968 (2021).

23. Saglam, A. S. & Chong, L. T. Protein–protein binding pathways and calculations of rate constants using fully-continuous, explicit-solvent simulations. *Chem. Sci.* **10**, 2360–2372 (2019).

24. Jorgensen, W. L., Chandrasekhar, J., Madura, J. D., Impey, R. W. & Klein, M. L. Comparison of simple potential functions for simulating liquid water. *J. Chem. Phys.* **79**, 926–935 (1983).

25. Case, D. A. *et al.* AMBER 2020. at (2020).

26. Crowley, M. F., Williamson, M. J. & Walker, R. C. CHAMBER: Comprehensive support for CHARMM force fields within the AMBER software. *Int. J. Quantum Chem.* **109**, 3767–3772 (2009).

27. Salomon-Ferrer, R., Götz, A. W., Poole, D., Le Grand, S. & Walker, R. C. Routine Microsecond Molecular Dynamics Simulations with AMBER on GPUs. 2. Explicit Solvent Particle Mesh Ewald. *J. Chem. Theory Comput.* **9**, 3878–3888 (2013).

28. Phillips, J. C. *et al.* Scalable molecular dynamics on CPU and GPU architectures with NAMD. *J. Chem. Phys.* **153**, 44130 (2020).

29. Loncharich, R. J., Brooks, B. R. & Pastor, R. W. Langevin dynamics of peptides: The frictional dependence of isomerization rates ofN-acetylalanyl-N'-methylamide. *Biopolymers* **32**, 523–535 (1992).

30. Åqvist, J., Wennerström, P., Nervall, M., Bjelic, S. & Brandsdal, B. O. Molecular dynamics simulations of water and biomolecules with a Monte Carlo constant pressure algorithm. *Chem. Phys. Lett.* **384**, 288–294 (2004).

31. Russo, J. D. *et al.* WESTPA 2.0: High-Performance Upgrades for Weighted Ensemble Simulations and Analysis of Longer-Timescale Applications. *J. Chem. Theory Comput.* **18**, 638–649 (2022).

32. Hsieh, C.-L. *et al.* Prefusion-stabilized SARS-CoV-2 S2-only antigen provides protection against SARS-CoV-2 challenge. *Nat. Commun.* **15**, 1553 (2024).

33. Torrillo, P. A., Bogetti, A. T. & Chong, L. T. A Minimal, Adaptive Binning Scheme for Weighted Ensemble Simulations. *J. Phys. Chem. A* **125**, 1642–1649 (2021).

34. Roe, D. R. & Cheatham, T. E. PTRAJ and CPPTRAJ: Software for Processing and Analysis of Molecular Dynamics Trajectory Data. *J. Chem. Theory Comput.* **9**, 3084–3095 (2013).

35. Suárez, E. *et al.* Simultaneous Computation of Dynamical and Equilibrium Information Using a Weighted Ensemble of Trajectories. *J. Chem. Theory Comput.* **10**, 2658–2667 (2014).

36. Bhatt, D., Zhang, B. W. & Zuckerman, D. M. Steady-state simulations using weighted ensemble path sampling. *J. Chem. Phys.* **133**, 14110 (2010).

37. Cohen, J. & Grayson, P. AutoIMD User's Guide. http://www.ks.uiuc.edu/Research/vmd/plugins/autoimd/ug/ (2011) doi:http://www.ks.uiuc.edu/Research/vmd/plugins/autoimd/ug/.

38. Bogetti, A. T. *et al.* A Suite of Advanced Tutorials for the WESTPA 2.0 Rare-Events Sampling Software [Article v2.0]. *Living J. Comput. Mol. Sci.* **5**, 1655–1655 (2022).

39. Efron, B. & Tibshirani, R. Bootstrap Methods for Standard Errors, Confidence Intervals,

and Other Measures of Statistical Accuracy. *Stat. Sci.* **1**, 54–75 (1986).

40. Michaud-Agrawal, N., Denning, E. J., Woolf, T. B. & Beckstein, O. MDAnalysis: A toolkit for the analysis of molecular dynamics simulations. *J. Comput. Chem.* **32**, 2319–2327 (2011).

41. Hunter, J. D. Matplotlib: A 2D Graphics Environment. *Comput. Sci. Eng.* **9**, 90–95 (2007).

42. Scheurer, M. *et al.* PyContact: Rapid, Customizable, and Visual Analysis of Noncovalent Interactions in MD Simulations. *Biophys. J.* **114**, 577–583 (2018).

43. Scheurer, M. *et al.* PyContact: Rapid, Customizable, and Visual Analysis of Noncovalent Interactions in MD Simulations. *Biophys. J.* **114**, 577–583 (2018).

44. Pedregosa, F. *et al.* Scikit-learn: Machine Learning in Python. *J. Mach. Learn. Res.* **12**, 2825–2830 (2011).

45. Waskom, M. seaborn: statistical data visualization. *J. Open Source Softw.* **6**, 3021 (2021).

46. Pettersen, E. F. *et al.* UCSF Chimera: A visualization system for exploratory research and analysis. *J. Comput. Chem.* **25**, 1605–1612 (2004).

47. Gapsys, V., Michielssens, S., Seeliger, D. & de Groot, B. L. pmx: Automated protein structure and topology generation for alchemical perturbations. *J. Comput. Chem.* **36**, 348–354 (2015).

48. Bauer, P., Hess, B. & Lindahl, E. GROMACS 2022 Manual. (2022) doi:10.5281/ZENODO.6103568.

49. Abraham, M. J. *et al.* GROMACS: High performance molecular simulations through multi-level parallelism from laptops to supercomputers. *SoftwareX* **1–2**, 19–25 (2015).

50. Goga, N., Rzepiela, A. J., de Vries, A. H., Marrink, S. J. & Berendsen, H. J. C. Efficient Algorithms for Langevin and DPD Dynamics. *J. Chem. Theory Comput.* **8**, 3637–3649 (2012).

51. Van Gunsteren, W. F. & Berendsen, H. J. C. A Leap-frog Algorithm for Stochastic Dynamics. *Mol. Simul.* **1**, 173–185 (1988).

52. Berendsen, H. J. C., Postma, J. P. M., Van Gunsteren, W. F., Dinola, A. & Haak, J. R. Molecular dynamics with coupling to an external bath. *J. Chem. Phys.* **81**, 3684–3690 (1984).

53. Hess, B. P-LINCS: A Parallel Linear Constraint Solver for Molecular Simulation. *J. Chem. Theory Comput.* **4**, 116–122 (2008).

54. Parrinello, M. & Rahman, A. Polymorphic transitions in single crystals: A new molecular dynamics method. *J. Appl. Phys.* **52**, 7182–7190 (1981).

55. Seeliger, D. & de Groot, B. L. Protein Thermostability Calculations Using Alchemical Free Energy Simulations. *Biophys. J.* **98**, 2309–2316 (2010).

56. Gapsys, V., Seeliger, D. & de Groot, B. L. New Soft-Core Potential Function for Molecular Dynamics Based Alchemical Free Energy Calculations. *J. Chem. Theory Comput.* **8**, 2373–2382 (2012).

57. Crooks, G. E. Entropy production fluctuation theorem and the nonequilibrium work relation for free energy differences. *Phys. Rev. E* **60**, 2721–2726 (1999).

58. Shirts, M. R., Bair, E., Hooker, G. & Pande, V. S. Equilibrium Free Energies from Nonequilibrium Measurements Using Maximum-Likelihood Methods. *Phys. Rev. Lett.* **91**, 140601 (2003).

59. Punjani, A., Rubinstein, J. L., Fleet, D. J. & Brubaker, M. A. cryoSPARC: algorithms for rapid unsupervised cryo-EM structure determination. *Nat. Methods* **14**, 290–296 (2017).
